# Identification of a Broadly Acting Inhibitor of the Alphavirus Non-Structural Protein 2 Helicase

**DOI:** 10.1101/2025.06.03.657755

**Authors:** John D. Sears, Durbadal Ojha, Sharon Taft-Benz, Paul Sylvester, Zachary Streblow, Amanda J. Gramm, Adam M. Drobish, Bose Muthu Ramalingam, Mohammad Anwar Hossain, Kacey Talbot, Wes Sanders, Che-Kang Chang, Isabella Law, Sabian A. Martinez, Jane E. Burdick, Renick Wiltshire, Nicholas A. Saba, Jack H. Poplarski, Rafael M. Couñago, Peter J. Brown, Ganesh S. Anand, Thomas E. Morrison, Jamie J. Arnold, Craig E. Cameron, Daniel N. Streblow, Timothy M. Willson, Mark T. Heise, Nathaniel J. Moorman

## Abstract

Alphaviruses are mosquito-borne viruses that have caused significant outbreaks in the 21^st^ century. Despite multiple recent outbreaks, there are no approved antiviral drugs to treat any alphavirus infection. Therefore, developing broadly acting antiviral drugs effective against multiple alphaviruses is necessary and could provide protection from both current and emerging alphavirus threats. A critical component of the alphavirus replication complex is non-structural protein 2 (nsP2), which is a multifunctional enzyme containing a helicase domain connected to a protease domain by a flexible linker. nsP2 functions as an ATP-dependent helicase, is conserved across the alphavirus genus, and is essential for virus replication, making it a promising target for development of alphavirus broad-acting antiviral drugs. Previous studies identified an enantioselective compound RA-0025298 that inhibited nsP2 ATPase activity and chikungunya virus CHIKV replication. Antiviral testing of RA-25298. against a diverse group of alphaviruses found broad activity except for Sindbis-like viruses. Using this information along with mutational profiling of virus passaged with RA-0025298 we identified the site of RA-0025298 action and confirmed the binding site via biophysical analyses. Finally, we found that the active enantiomer of RA-0025298 (SGC-NSP2hel-1) reduced viral loads in vivo and protected mice from tissue damage and disease caused by CHIKV infection. These findings further describe the mechanism of action of a first-in-class nsP2 helicase inhibitor with the potential for development as a broad spectrum drug for treating or preventing disease caused by current and emerging alphaviruses.

**One Sentence Summary:** This study describes the mechanism of action and in vivo efficacy of a first in class broadly acting inhibitor of alphavirus nsP2 helicase activity.

## Introduction

Alphaviruses are mosquito-borne viruses that have emerged in the 21^st^ century to cause significant outbreaks across the globe. Alphaviruses are broadly classified into two groups, arthritogenic and encephalitic, based on the disease they cause (*1*, *2*). Arthritogenic alphaviruses including Chikungunya virus (CHIKV) have caused multiple widespread outbreaks of severe acute and chronic arthralgia (*3*, *4*). Encephalitic alphaviruses like Eastern equine encephalitis virus cause sporadic outbreaks of viral encephalitis with high morbidity and mortality. Previous efforts to weaponize encephalitic alphaviruses highlight their potential as bioterrorism weapons (*5–7*). In addition, emerging alphaviruses such as Mayaro virus (MAYV) are poised for rapid expansion into immunologically naïve human populations as the range of their mosquito vectors continues to increase (*8*). Despite the clear and present danger presented by alphaviruses, there are currently no FDA approved drugs to treat any alphavirus infection and limited vaccine availability (*9*, *10*). The development of medical countermeasures broadly active against alphaviruses is thus critically important for treating and preventing disease caused by current and emerging alphaviruses.

The alphaviruses non-structural proteins are especially promising targets for antiviral therapies due to their high sequence conservation and essential roles in virus replication (*11–13*). Alphaviruses are positive sense single stranded RNA viruses with two open reading frames. The full-length genomic RNA (gRNA) encoding the four non-structural proteins (nsP1-4) that together comprise the viral replicase, is translated immediately upon host cell entry. The replicase transcribes a sub-genomic RNA (sgRNA) encoding the viral structural proteins, as well as negative strand templates for the synthesis of new positive strand genomes (*14*, *15*). In addition to their functions in RNA synthesis, the nsPs also play key roles in remodeling infected cells to support replication and subverting the host innate response to infection (*16–18*). Thus, small molecule drugs targeting nsPs could have the dual benefit of directly inhibiting virus replication while simultaneously enhancing the host antiviral immune response.

Of the non-structural proteins, nsP2 protein is an especially promising drug target because of its multiple essential roles in alphavirus replication (*19*). nsP2 encodes an amino terminal helicase domain and carboxy terminal protease domain linked by a flexible region (*20*, *21*). Both the protease and helicase activities are essential for virus replication. The protease domain processes the viral non-structural polyprotein encoded by gRNA into the individual subunits of the replicase (*14*, *22*), while the helicase domain is required for capping of viral RNA, viral RNA synthesis, and the inhibition of host cell transcription and translation (*23–25*). The high sequence conservation of nsP2 (>50% amino acid identity across all alphaviruses, Fig.S1A) makes it a prime target for development of broadly acting antiviral drugs for both current and newly emerging alphavirus threats.

While several groups have developed small molecule nsP2 protease inhibitors (*26–30*), the identification of direct acting nsP2 helicase inhibitors has proven more challenging. The nsP2 helicase domain (nsP2hel) is a member of the SF1B helicase family and contains a RecA1 domain with a Walker A and B motif needed for its NTPase activity and a RecA2 domain critical for RNA binding (*31*). As with other SF1B helicases the NTPase activity of nsP2hel drives its translocation and RNA unwindase activity, which is required for RNA synthesis and viral replication (*32–35*). However, the nsP2hel ATP and RNA binding sites are structurally similar to those in cellular helicases, raising the risk of off target effects (*36*, *37*). Helicases also undergo dynamic conformational changes throughout the enzyme cycle, making the identification of well-defined small molecule binding sites for allosteric inhibition challenging (*38*).

Despite these challenges, our previous efforts identified novel inhibitors of CHIKV nsP2 ATPase activity (*39*). Hit optimization resulted in the discovery of RA-25298, a specific, non-competitive inhibitor of nsP2 ATPase and unwindase activity that potently inhibits CHIKV replication in vitro (*40*). In this study we show that RA-25298 inhibits viral replication by blocking RNA synthesis and inhibits the replication of multiple viruses in the alphavirus family. Using mutational profiling, comparative analysis of nsP2 from different alphaviruses, and biophysical studies we identified a potential binding site and allosteric mechanism of action for this compound. Importantly, SGC-NSP2hel-1, the active R-enantiomer of RA-25298 (*41*), reduced virus replication and limited virus-induced pathology in a preclinical small animal model of CHIKV disease. These data identify SGC-NSP2hel-1 as a novel, specific inhibitor of the nsP2 helicase with broad spectrum activity against multiple alphaviruses and provides new insights into drug discovery and development strategies useful for targeting additional SF1B helicases in other viral and human diseases.

## Results

### RA-25298 inhibits virus replication by inhibiting viral RNA synthesis

We previously identified RA-25298 as a potent and specific small molecule inhibitor of the CHIKV nsP2 helicase activity (*40*). To understand how RA-25298 inhibits CHIKV replication, we used a series of cell-based assays to measure its effect on helicase action and virus replication. We found that RA-25298 inhibits CHIKV virus replication in a dose-dependent manner (Fig. 1A). When added prior to infection, RA-25298 decreased CHIKV non-structural and structural protein synthesis (Fig. 1B) and inhibited the accumulation of viral RNA (Fig. 1C). To define the temporal efficacy of RA-25298 throughout the viral lifecycle, we added RA-25298 at different times after infection and measured its effect on the accumulation of cell-free infectious virus. Addition of RA-25298 as late as 12 h post infection significantly reduced virus replication demonstrating the therapeutic potential of RA-25298 (Fig. 1D). These data show that RA-25298 inhibits CHIKV replication by inhibiting the synthesis of viral RNA leading to a decrease in protein expression, consistent with the known roles of nsP2 helicase activity in the viral replicative cycle.

**Figure 1.**
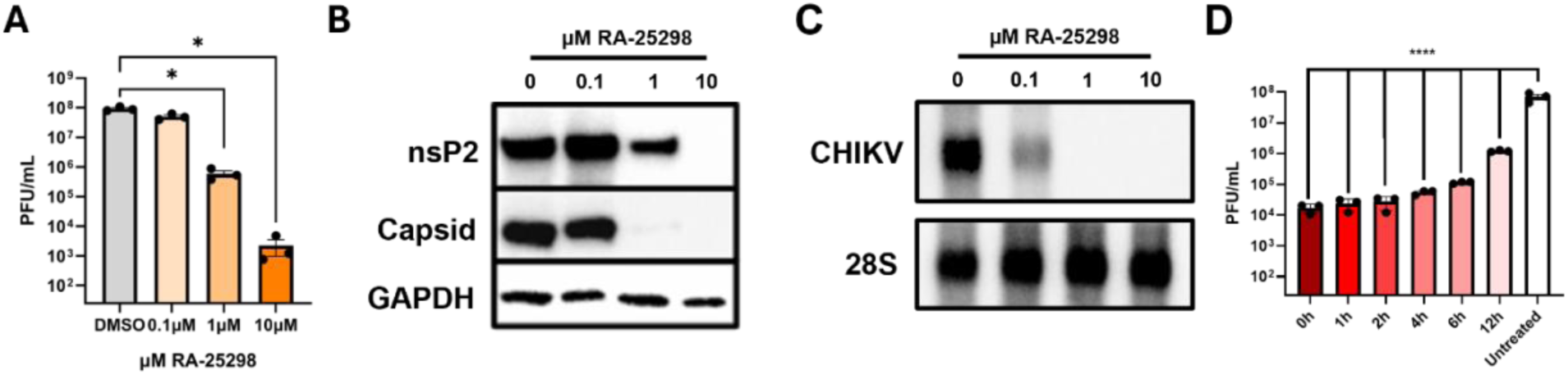
RA-25298 inhibits CHIKV replication by inhibiting viral RNA synthesis and protein expression. (A) MRC-5 fibroblasts were treated with the indicated concentrations of RA-25298 for one hour before infection with CHIKV (MOI=0.1). Samples were collected 24 hours after infection and levels of infectious virus were measured by plaque assay. (B) Cells were treated and infected as in (A), and the synthesis of viral non-structural (nsP2) and structural (Capsid) proteins and (C) viral RNA was measured by Western blot or Northern blot, respectively. (D) A time of addition study was performed by adding 1.25uM RA-25298 at different times after CHIKV. Levels of infectious virus were measured at 24 hours after infection (n=3).

### RA-25298 is a broadly acting inhibitor of alphavirus replication except for Sindbis-like Viruses

To measure the breadth of RA-25298 activity, we tested its antiviral efficacy against a wide range of alphaviruses. Based on the amino acid sequence of nsP2hel, the alphavirus family can be divided in to three groups: Old World, New World (EEEV and VEEV like viruses), and a distinct subgroup of Old World viruses closely related to Sindbis virus (SINV, Fig. 2A, Fig. S1A). RA-25298 inhibited the replication of representative Old and New World alphaviruses, however SINV-like viruses including SINV, Whataroa and Aura virus were resistant to inhibition by RA-25298, even at concentrations >200 fold higher than the EC_50_ for CHIKV (Fig. 2B-E, Fig.S2 A-M). Similar results were observed in biochemical assays measuring nsP2 ATPase and RNA unwindase activity, where RA-25298 showed potent inhibition of Old (CHIKV) and New (VEEV) World nsP2 orthologs, while SINV nsP2 was resistant to inhibition by RA-25298 (Fig.2F,G). These data confirm that RA-25298 is a broadly acting inhibitor of alphavirus nsP2 ATPase and unwindase activity and virus replication. The resistance of SINV nsP2 to inhibition suggests that SINV-like nsP2 sequences that differ with nsP2 from other alphaviruses are likely critical for RA-25298 activity.

**Figure 2.**
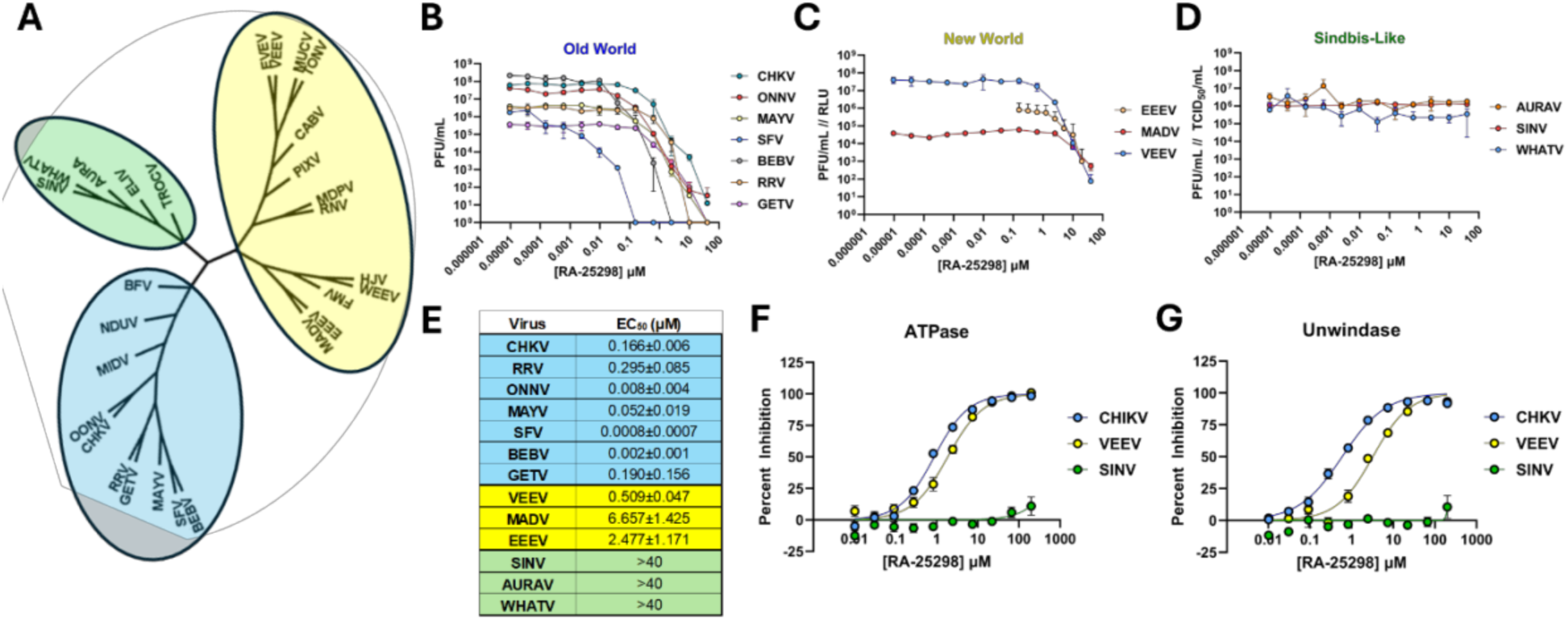
RA-25298 is broadly acting inhibitor of alphavirus replication. (**A**) Phylogenetic analysis of the nsP2 helicase domain divides the family into three distinct clades. (B-D) Cells were infected as in Figure 1, and the effect of RA-25298 on the replication of Old World (CHIKV, RRV, ONNV, MAYV), New world (EEEV, VEEV, MADV), or Sindbis-like (SINV) alphaviruses. (E) Table summarizing the 50% effective antiviral concentration (EC50) of RA-25298 on a panel of alphaviruses. (F, G) Dose response curves showing the effect of RA-25298 on the ATPase (F) or unwindase (G) activity of purified nsP2 from CHIKV, VEEV or SINV (N=3,MOI=0.1).

### RA-25298 targets an allosteric site proximal to the nsP2 RecA1:RecA2 interface but distinct from the ATP binding site

To define the potential site of action for RA-25298 in nsP2 we serially passaged CHIKV in the presence of the inhibitor. Passaging three separate lineages of CHIKV in the presence of the EC_99_ of RA-25298 led to an increase in viral titers over the course of six passages (Fig. 3A). All lineages were fully resistant to inhibition by RA-25298 (10µM) by the sixth passage, while controls virus passaged in the presence of the vehicle control (DMSO) remained susceptible to inhibition (Fig. 3B). Mutations in each lineage were identified by long-read sequencing, revealing multiple mutations associated with resistance. One mutation, F185L occurred at the highest frequency in each of the three lineages (Fig. 3C, Fig.S3A-C), suggesting an important role for this residue in inhibition of nsP2 helicase activity by RA-25298.

**Figure 3.**
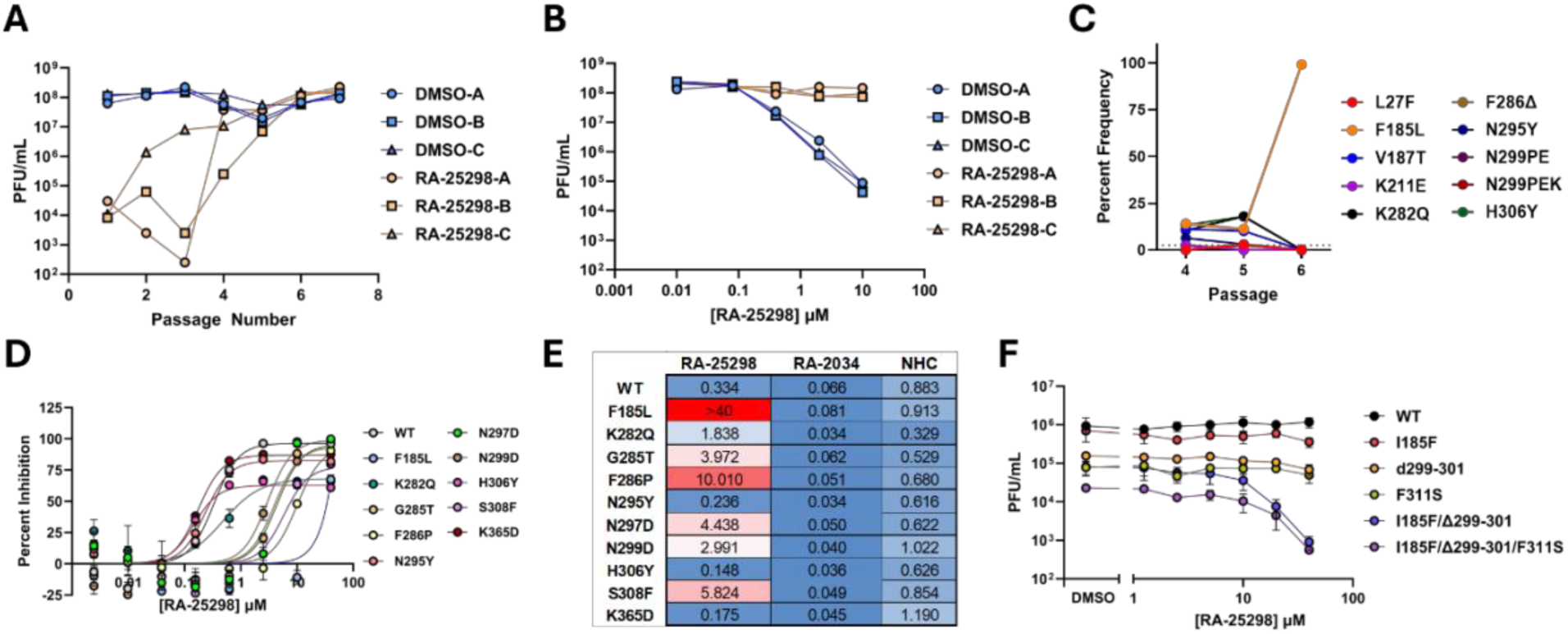
RA-25298 interacts with an allosteric target site at the junction of the RecA1 and RecA2 domain. (A) Chikungunya virus (181/25) was serially passaged in the presence of RA-25298 or DSMO. Viral titers were by plaque assay after each passage to monitor for RA-25298 resistance. (B) After the 6^th^ passage each independent passage lineage was tested in a titer reduction assay with a 4-fold 6 point dose response of RA-25298 starting at 10uM. (C) The abundance of mutations with >5% penetrance in viruses passaged in the presence of RA-25298 was determined by next generation sequencing. A representative plot shown. (D) Mutations correlating with resistance were made in a CHIKV infectious clone containing a nano-luciferase reporter gene. Each mutant was tested for RA-25298 sensitivity infected cells over an 8-point 4-fold dose response starting at 40uM. (E) Table listing the EC50 values of each mutant for RA-25298, an alphavirus protease inhibitor (RA-2034) or an inhibitor of the alphavirus replicase (NHC). (F) Titer reduction assay showing the effect of RA-25298 on the replication of a panel of SINV mutants (G100 strain). (G) Docking model of RA-25298 displayed overlayed onto a three dimensional model of nsP2. In the left panel, nsP2 shading reflects hydrophobicity. In the center and rightmost panel, nsP2 residues associated with resistance are shown in red, and proximal residues in the Walker A and Walker B motifs are shown in purple and yellow, respectively. Dashed lines represent potential interaction sites between nsP2 and RA-25298.

Each mutation correlating with resistance was individually introduced into a previously described (*42*) infectious clone of the CHIKV 181/25 vaccine strain containing a nanoLuciferase (nLuc) reporter gene inserted between the capsid and E3 genes, and then tested for inhibition by RA-25298. Three mutants (F185L, N299D, and S308F) had antiviral and enzymatic EC_50_ values > 2.5µM, a significant increase as compared to wild type virus, indicating resistance to the inhibitor. An independent study identified a different cluster of amino acids that could be potential target sites for small molecule inhibition of the nsP2 helicase. Mutations in three of these amino acids (G285T, F286P, and N297D) conferred similar resistance to RA-25298 in both antiviral and enzymatic assays(Fig 3D, Fig.S4 A,FigS5 A). Orthologous mutations made in an EEEV replicon (Y185L,G285T, and F286P) were also resistant to inhibition by RA-25298 (Fig.S5B). All CHIKV mutants conveyed resistance specifically to RA-25298, as they remained fully susceptible to the nsP2 protease inhibitor RA-2034 (*26–28*)) and the nsP4 polymerase inhibitor β-d-N4-hydroxycytidine (NHC) (*43*) (Fig. 3E, Fig S5A). Of note, each of the mutants had ATPase, unwindase, and RNA binding activity comparable to wild type nsP2, with the exception of G285T and F286P, which increase and decrease unwindase activity, respectively (Fig.S4 A-G). This data is consistent with previous studies showing that this region of the protein regulates ATPase and unwindase activity (*31*).

When mapped onto the three-dimensional structure of nsP2hel, the resistant mutations cluster around a pocket opposite to the ATP binding site but proximal to an exposed residue of the Walker B motif (F255). Interestingly, SINV and SINV-like viruses differ significantly from Old and New World alphaviruses in this region. SINV-like viruses contain a three amino acid insertion (P299,E300,K301) not present in Old or New World alphaviruses that forms a small alpha helix that occludes the pocket (Fig. S6A). Amino acids surrounding this pocket that differ between CHIKV and SINV acquired mutations during serial passage with RA-25298 (Fig. S6A-C). Several of the CHIKV RA-25298 resistance mutations resulted in changes to the corresponding amino acid naturally found in SINV, including the insertion of the three amino acid alpha helix and a change at amino acid 185 which is similar to the CHIKV F185L resistance mutation (Fig. S3A-C).

The above data suggested that the nature of the amino acids surrounding the pocket are key determinants of RA-25298 inhibition. To further test this hypothesis, we mutated SINV amino acids in this region to the corresponding amino acids found in CHIKV, both alone and in combination. No single mutation could confer SINV susceptibility to RA-25298. However, the deletion of the three amino acid alpha helix (Δ299-301) in combination with the I185F and F311S mutations significantly increased SINV susceptibility to RA-25298 in both antiviral and enzymatic assays (Fig. 3F, Fig. S6D,E). Interestingly, the SINV Δ299-301 mutation alone significantly reduced unwindase and ATPase Vmax. However, including the additional mutations that enhanced sensitivity to RA-25298 restored ATPase and unwindase Vmax to wild type levels but these additions increase the ATPase Km indicating that interactions between these residues may be important for SINV ATP affinity (Fig. S6F-H).

### SGC-nsP2hel-1 forms a stable complex with nsP2 and ATP

To further understand the mechanism of nsP2 ATPase inhibition by RA-25298 analogs we performed biophysical analyses of the full length nsP2 protein in the presence of ATP and non-hydrolysable ATP analogs. RA-25298 is a racemic mixture of two enantiomers (*41*). We previously demonstrated that only the R-enantiomer SGC-NSP2hel-1 (aka RA-0188293) maintains potent nsP2 helicase inhibition while its S-enantiomer SGC-NSP2hel-1N (aka RA-0188294) is >100-fold less active (*41*). Differential scanning fluorimetry (DSF) assays with CHIKV nsP2 in the presence of RA-25298 or its active R-enantiomer SGC-NSP2hel-1 found that both compounds increased nsP2 thermal stability, while the inactive S-enantiomer did not (Fig. 4A). Because SGC-NSP2hel-1 is a non-competitive inhibitor of nsP2 ATPase activity we tested its effect on nsP2 stability in the presence of ATP and non-hydrolysable ATP analogs. Slow hydrolyzing analogs such as ATPγS led to an increase in CHIKV nsP2 stability by themselves. However, the combination of ATP or the ATP analogs with SGC-NSP2hel-1 resulted in a large thermal shift (Fig. 4B). The increase in stability was not observed with CHIKV nsP2 protein containing the F185L resistance mutation or with SINV nsP2 protein, suggesting that the lack of inhibition of the different RA-25298-resistant genotypes is driven by an inability of SGC-NSP2hel-1 to interact with and stabilize nsP2 (Figure 4C,D). This data indicates that nsP2 forms a stable complex with SGC-NSP2hel-1 and ATP.

**Figure 4.**
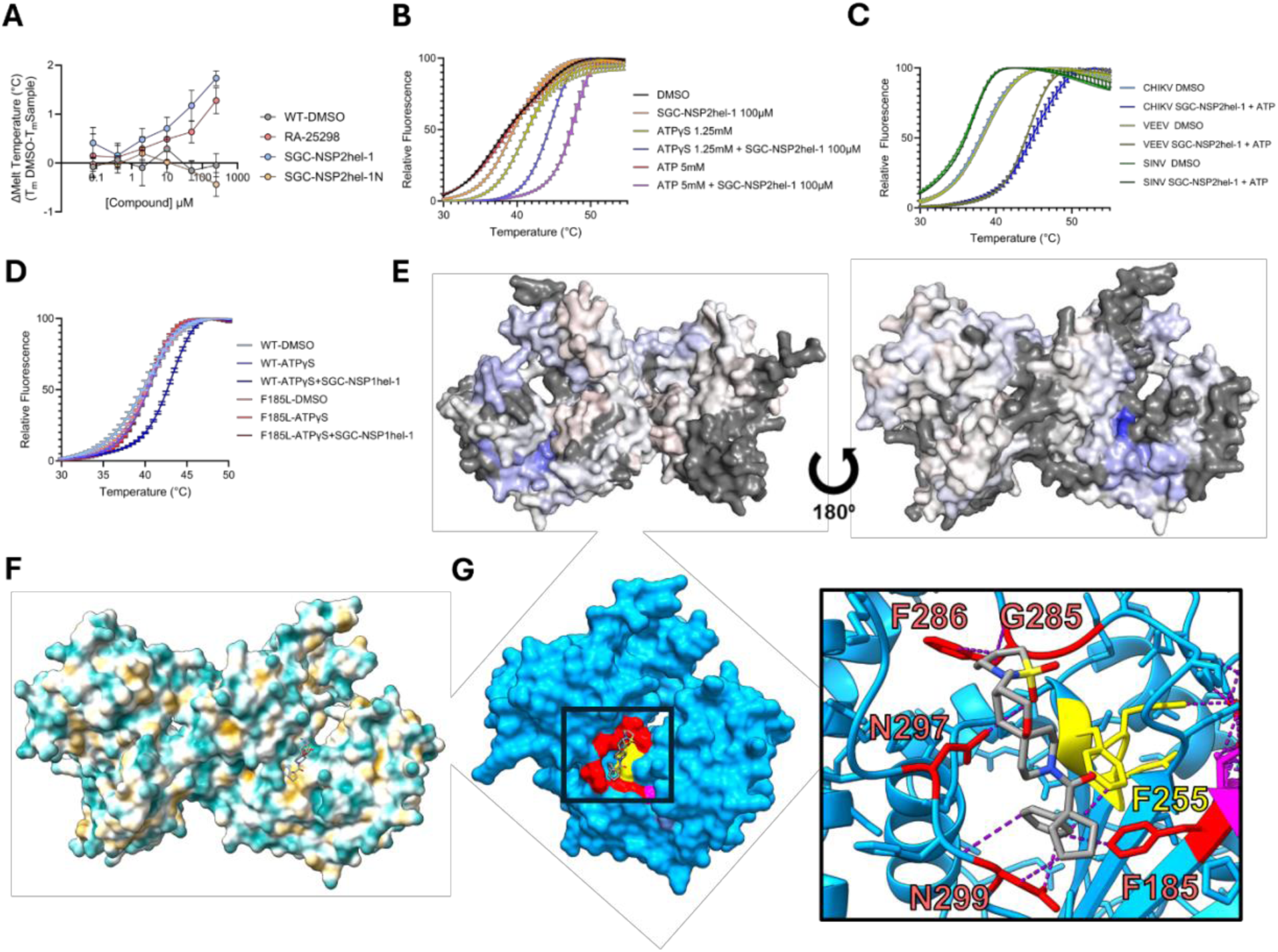
ATP cofactor binding stabilizes SGC-NSP2hel-1interaction with nsP2. (A) Differential scanning fluorimetry (DSF) assays was performed using CHIKV nsP2 with a 6 point 4 fold dose response of RA-25298, RA-0188293, or RA-0188294. Change in melt temperature as determined by Boltzmann sigmoidal curve was measured by comparing melt temperature with compound to DMSO. (B-D) DSF assay measuring relative fluorescence was performed with CHIKV, VEEV, SINV, or CHIKV F185L nsP2 in the presence of different combinations of 100µM SGC-NSP2hel-1and 5mM ATP or 1.25mM ATPγS. (E) ATP hydrolysis by CHIKV in a single cycle reaction was measured with p32 labeled ATP in the presence of RA-0188293. (F) Phosphate release in a single cycle reaction was measured by MDCC assay. (G) CHIKV nsP2 surface exposure with ATPγS and SGC-NSP2hel-1compared to apo was measured by HDX-MS and difference plotted by heat map with blue indicating regions more protected in the presence of ATPγS and SGC-NSP2hel-1and red more exposed.

Because of the dynamic nature of SF1B helicases, structural studies to capture intermediate complexes can be challenging. Therefore, to determine of how the nsP2, SGC-NSP2hel-1, and ATP form a stable complex we performed comparative HDX-MS with all combinations of nsP2, ATPγS, and SGC-NSP2hel-1 (Table S1). 123 pepsin fragments were observed, resulting in a coverage of 77.9% of nsP2 (Figure S7A). HDX-MS was measured at deuterium exchange times (D_ex_ = 1 and 10 min). Interestingly, no differences in deuterium uptake were seen between nsP2 alone and nsP2 with ATPγS (Fig. S8A,Fig.S9A). Comparing nsP2 alone to nsP2 in the presence of SGC-NSP2hel-1 identified one peptide (185–201) showing significant difference at D_ex_ = 10 min (Fig. S9B. A more pronounced difference was found when comparing nsP2 alone to nsP2 in the presence of both SGC-NSP2hel-1 and ATPγS with deuterium exchange significantly altered for 3 peptides (221-230, 250-255, and 239-302) at D_ex_ = 1 min and 5 peptides (64-82, 182-201, 185-If201, 250-255, and 293-301) at D_ex_ = 10 min (Fig.S9C-E) Interestingly these peptides contain the amino acids most frequently mutated in resistance passaging experiments (Fig.3E). The largest observed change in protection comes from the peptides (182-201 and 185-201) that include F185. (Fig.4E), consistent with mutational profiling data showing that an F185L mutation in the CHIKV nsP2 helicase domain confers resistance to inhibition by SGC-NSP2hel-1.

Data from passaging and mutagenesis experiments, comparative analysis of the effect of RA-25298 on diverse alphaviruses, as well as DSF and HDX-MS data were integrated into a docking model and overlayed on a model of nsP2 structure colored by hydrophobicity (Fig. 4F). In this model, the RA-25298 amide group interacts with a hydrophobic pocket between N299 and F185L, with the spirocyclic core positioned in a cleft between the Walker B motif (D252-F255) and N295 and N297 and the sulfonamide group positioned towards G285 and F286 (Fig. 4G).

### SGC-NSP2hel-1 reduces viral replication and disease in a preclinical model of CHIKV infection

SGC-NSP2hel-1 is a potent and specific nsP2 inhibitor active against multiple alphaviruses in infected cells with very little impact on cell viability with a CC_50_ >250µM and Selectivity Index of >4000. Initial pharmacokinetic profiling of SGC-NSP2hel-1 showed high clearance in mice due to rapid first pass metabolism. However, when co-administered with the p450 inhibitor 1-ABT, oral dosing of SGC-NSP2hel-1 at 10 mg/kg resulted in plasma concentrations exceeding the CHIKV EC_90_ (Fig. S10B,C) for 3h supporting its testing in a preclinical model of CHIKV replication and pathogenesis. Mice were dosed orally twice daily with 30mg/kg SGC-NSP2hel-1 and once daily with 100mg/kg 1-ABT (Fig. 5A), beginning at 12 hours prior to infection with CHIKV. Following infection, mice treated with SGC-NSP2hel-1 or the vehicle control maintained body weight over the course of the study (Fig. S11A). In this model, CHIKV replication induces significant swelling of the ipsilateral foot and ankle, in line with the severe inflammation of the joints observed in people with acute CHIKV disease (*2*, *44*, *45*). SGC-NSP2hel-1 treatment significantly reduced footpad swelling compared to the vehicle control (Figure 5B) and significantly reduced virus replication in the ipsilateral foot and serum (Fig. 5C-E, Fig. S8 B-F). SGC-NSP2hel-1 treatment also significantly decreased virus-induced pathology compared to animals treated with vehicle control, with decreased inflammatory cell recruitment and reduced myositis and muscle damage (Fig. 5F-I, Fig. S8 G,H). These data reveal SGC-NSP2hel-1 to be a promising new small molecule for treating CHIKV disease.

**Figure 5.**
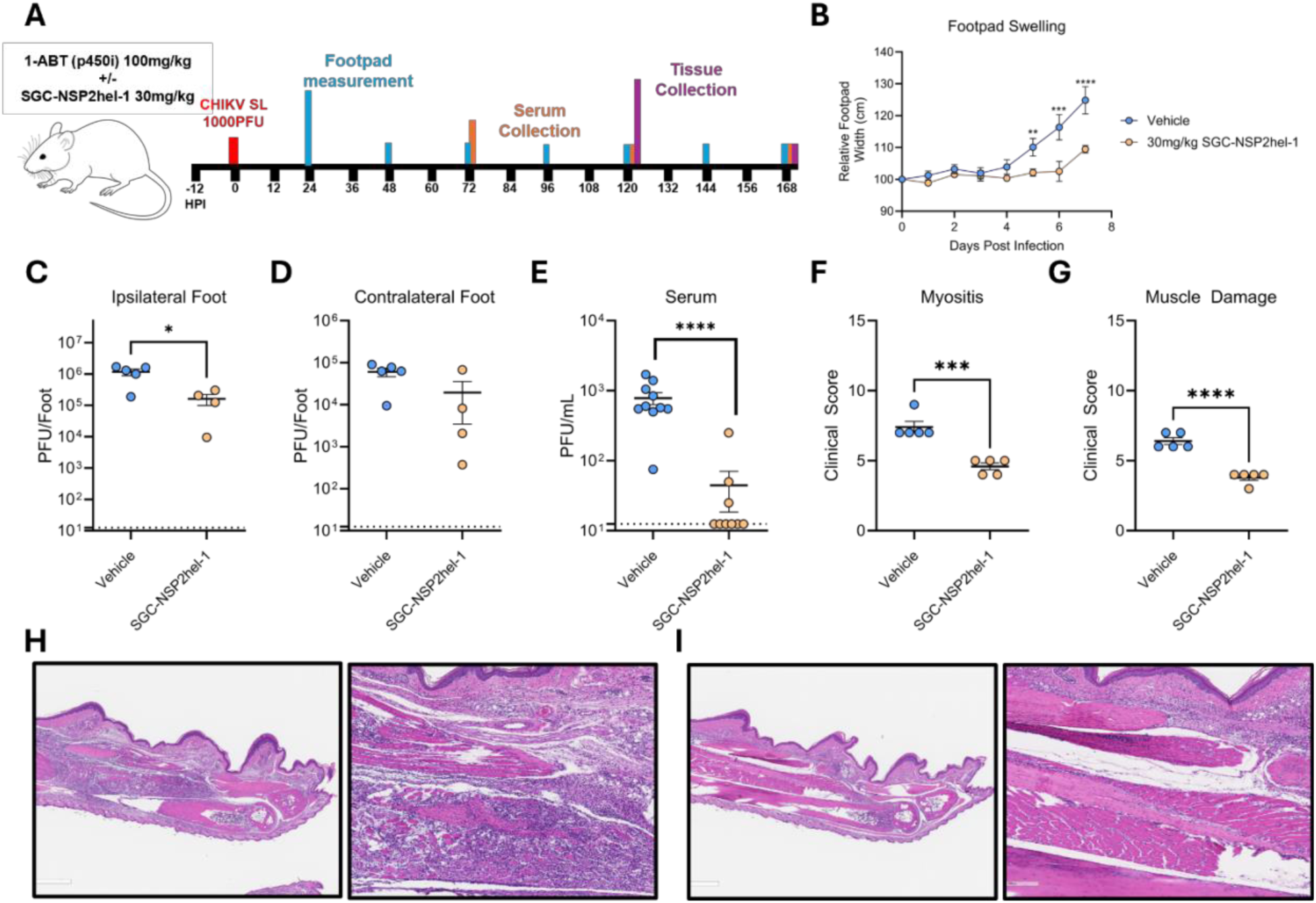
RA-001888293 displays efficacy against replication and pathology in mouse model of chikungunya virus infection. (A) Experimental design of in vivo efficacy study measuring the effect of prophylactic of co-dosing SGC-NSP2hel-1(30mg/kg BID) and a cytochrome P450 inhibitor (1-ABT 100mg/kg once daily) or a vehicle control on CHIKV replication and disease. (B) Relative footpad swelling throughout the first 7 days of infection in SGC-NSP2hel-1or vehicle treated mice. (C-E) Viral titers were measured in the indicated tissues at 5 days after infection except for serum, which was measured at 3 days post infection. (F-H) Feet from infected mice were collected at 7 days after infection. H&E stained sections were assessed for pathology in a blinded analysis to quantify (F) and muscle damage (G).

## Discussion

Here we define the antiviral mechanism of action of RA-25298, a first-in-class broad spectrum allosteric inhibitor of the alphavirus nsP2 helicase. RA-25298 significantly reduces viral RNA synthesis, inhibits virus replication when added before or after CHIKV infection and inhibits the replication of a wide range of alphaviruses in vitro demonstrating its potential as a broad spectrum prophylactic and therapeutic drug to combat alphavirus disease. While RA-25298 has antiviral efficacy against most New and Old World alphaviruses, the distinctive SINV-like subgroup was resistant. Passaging CHIKV in the presence of RA-25298 resulted in resistance mutations mapping to a region of the helicase domain on the opposite face from the ATPase binding pocket. Interestingly, this binding site is adjacent to an alpha helical insert found in the RA-25298 resistant Sindbis subfamily and not in the other alphaviruses that are sensitive to RA-25298 inhibition. Mutagenesis, biochemical, and biophysical studies identified key amino acids in this region conferring resistance or sensitivity to RA-25298 in assays of virus replication, ATPase, and RNA unwindase assays, and DSF assays using purified nsP2 protein. HDX-MS analysis displayed significant protection of this region in the presence of SGC-NSP2hel-1 and ATPγS providing further support for this site as a key location to SGC-NSP2hel-1 Computational docking studies revealed that binding of RA-25298 and its active R-enantiomer SGC-NSP2hel-1 maps to a hydrophobic pocket composed of exposed residues of the Walker B motif. We demonstrated that SGC-NSP2hel-1 co-dosed with a P450 inhibitor was protective in a preclinical mouse model of CHIKV disease, significantly reducing virus titer and virus-induced pathogenesis. These data support further preclinical development of SGC-NSP2hel-1 for treating alphavirus disease and reveal features of allosteric inhibition that may be useful for targeting other RNA helicases involved in human diseases.

There are currently no approved antiviral drugs to treat or prevent any alphavirus infection. The identification of RA-25298 and its enantiomer SGC-NSP2hel-1 is an important first step in addressing this urgent unmet need. Typically, RNA helicases are considered to have low tractability as antiviral targets, in part because their dynamic motor function has not been conducive to identification of high affinity inhibitors (*38*, *46*).We previously reported that SGC-NSP2hel-1 specifically inhibits alphavirus nsP2, with limited activity against a panel of human RNA helicases (*41*). SGC-NSP2hel-1 has a high selectivity index in vitro and showed no gross toxicity in animals at doses above those required for antiviral efficacy in vivo. These data validate nsP2 helicase activity as a promising target for direct-acting antiviral drugs and support further preclinical development of SGC-NSP2hel-1 as a potent, non-toxic, broad spectrum inhibitor of alphavirus replication and disease. Furthermore, the availability of a highly potent and specific nsP2 helicase inhibitor provides a potent tool for defining how nsP2 NTPase and unwindase activity contribute to distinct stages of viral replicative cycle.

The specificity and lack of toxicity of RA-25298 and SGC-NSP2hel-1 likely reflects the allosteric mechanism of helicase inhibition. Rather than targeting the ATP or RNA binding sites, mutational profiling together with biochemical and biophysical data indicate that RA-25298 binds a hydrophobic pocket opposite the Walker B motif at the intersection of the RecA1 and RecA2 domains. While this region is largely conserved across alphavirus nsP2 orthologs, it differs sufficiently from cellular host RNA helicases to allow for specific binding and inhibition. This hypothesis is consistent with our data showing differences in RA-25298 susceptibility between subgroups of alphaviruses. While most alphaviruses are sensitive to RA-25298 inhibition, the SINV-like subgroup are resistant. Viruses in this subgroup contain a three-amino acid insertion at the predicted RA-25298 binding site which creates a small helix in the RecA2 domain, which would occlude inhibitor binding. Deletion of this loop resulted in a decrease in enzymatic activity in SINV nsP2; however, introducing an additional mutation (I185F) in the RecA1 domain to match the corresponding residue found in CHIKV and VEEV restored activity. These findings emphasize the critical role of RecA1–RecA2 domain interactions in regulating enzymatic function, and the ability of small molecules targeting this region to potently and specifically inhibit an RNA helicase in an allosteric manner. Other SF1B helicases contain a similar pocket at the RecA1:RecA2 interface, though the proximal amino acid sequence varies. While further studies with additional viral and human helicases are needed, our results suggest that targeting this region with small molecules may provide a common strategy for developing RNA helicases inhibitors for use in treating viral and human diseases.

Together our results suggest a model where SGC-NSP2hel-1 first binds to the nsP2 helicase domain in its ATP-free, open conformation. Mutational profiling and in silico docking suggest SGC-NSP2hel-1 binds in a hydrophobic pocket at the junction of the RecA1 and RecA2 domains between N299 and F185 without affecting RNA binding. ATP binding by nsP2 induces a conformational shift that brings its Walker A and B motifs in the RecA1 and RecA2 domains, respectively, into close proximity (*47*). Rather than disrupting SGC-NSP2hel-1 binding, this conformational shift appears to strengthen compound binding. The presence of SGC-NSP2hel-1 in the pocket prevents residues in the Walker B motif from reaching close enough proximity to hydrolyze ATP bound to the Walker A motif. As a result, the nsP2 helicase domain ‘locks’ in the closed conformation in the presence of SGC-NSP2hel-1, preventing further rounds of helicase action. While this model is consistent with our data, additional experiments will be needed to confirm the specific SGC-NSP2hel-1 binding pose.

Importantly, SGC-NSP2hel-1 inhibited virus replication and prevented disease in a preclinical mouse model of CHIKV infection. Prophylactic SGC-NSP2hel-1 treatment decreased CHIKV replication in tissues and serum while also preventing footpad swelling, myositis, and virus-induced muscle damage. These data further validate the nsP2 helicase as a therapeutic target for direct acting antiviral drugs and show that SGC-NSP2hel-1 is both orally bioavailable and distributed to sites of CHIKV disease. Given its potent in vitro antiviral efficacy, optimized SGC-NSP2hel-1 analogs with increased viral breadth and decreased cytochrome P450 metabolism are likely to be even more effective in treatment of alphavirus disease. The breadth of in vitro antiviral activity also suggests that SGC-NSP2hel-1 may be similarly efficacious at preventing disease caused by additional alphaviruses, providing a string rationale for further efforts to optimize SGC-NSP2hel-1 towards clinical use.

## Methods

### ADP-Glo ATPase Assay

CHIKV nsP2 ATPase activity was tested using the Promega ADP-glo assay kit in a 384-well format. 1 nM Purified CHIKV nsP2 was incubated with compound diluted in an 8-point 3-fold dose response starting at 250 µM for 30 minutes prior to addition of 30 µM ATP was added and incubated for another 1 hour at room temperature. 2 µl of ADP-Glo reagent (Promega) was added to the reaction and incubated for 40 minutes followed by the addition of Kinase detection reagent, a 30 minute incubation and reading the assay plate for luminescence using Tecan Spark.

### Unwindase Assay

CHIKV unwindase activity was measured using a FRET based duplex substrate with a Cy5 labeled RNA (5’-Cy5CGCUGAUGUCGCCUGG-3’) coupled to a BHQ labelled RNA (5’-AAAAAAAAAAAAACCAGGCGACATCAGCG-3’). 10nM CHIKV nsP2 in the presence of 125 nM of unlabeled competitor strand (5’-CCAGGCGACAUCAGCG-3’) was incubated with compound diluted in an 8-point 3-fold dose response starting at 250 µM for 30 minutes prior to addition of 25 nM of the FRET substrate. Fluorescence was read at 620 nm excitation and 685 nm emission with a manual gain of 180 on Tecan spark 30 minutes following the addition of the substrate.

### Reporter Virus Antiviral Assay

Compound antiviral activity was measured using a previously described nanoluciferase reporter assay using 181/25 CHIKV with nanoluciferase inserted behind the capsid gene (2). In brief, MRC-5 fibroblasts were treated with an 8 point 4 fold dose response of compound starting at 40µM for 1 hour prior to infection with 181/25 nanoluciferase capsid virus (MOI=0.1) and virus replication was determined by measuring nanoluciferase production using Nano-Glo® Luciferase Assay System (Promega) and read on a GloMax Navigator (Promega). Percent inhibition of the compound was calculated by comparing relative luciferase units of experimental wells to DMSO control wells.

### Thermal Shift Assay

Following a previously described protocol using SYBR Orange (*48*)5mM nsP2 or nsP2 helicase domain were incubated with treatment for 30 minutes in buffer optimized for CHIKV nsP2 (200mM NaCl,50mM HEPES pH 7.8). Samples were then run on a melt curve ranging from 20°C to 95°C 1c/min (Bio-Rad). Relative fluorescence was calculated by normalizing each curve with its lowest value and dividing each point by its maximum value and multiplying by 100. Melt temperatures were determined from relative fluorescence using a Boltzmann sigmoidal curve.

### RNA Binding

dsRNA substrates were produced by annealing RNAs 5’-UUUUUUUUUUUUUUUUUUUUGCCGCCCGGUUUUUUUUUU-3’, and 5’-FITC-CCGGGCGGCUUUUUUUUUU-3’ in 50 mM NaCl, 10 mM Tris-HCl, and 1 mM EDTA in a Progene Thermocycler (Techne) with a molar ratio of 1:1.4 for labeled:unlabeled strands. Annealing reaction mixtures were heated to 90 °C for 1 min and slowly cooled (5 °C/min) to 10 °C. Increasing concentrations of nsP2 were mixed with 1 nM of fluorescein-labeled dsRNA substrate in a binding buffer containing 25 mM HEPES pH 7.5, 1 mM MgAcetate, 1 mM TCEP, and 100 mM KGlutamate. Reactions were incubated at room temperature for 10 minutes in a 384-well non-binding plate and then read with a dual wavelength 485/20 485/20 fluorescence polarization cube using a Biotek Synergy H1 plate reader. Data from protein titration experiments were fit to a hyperbola to calculate dissociation constants.

### Passaging Experiment

MRC-5 fibroblasts were infected with WT CHIKV (181/25) at an MOI of 0.01 treated either with the antiviral EC_99_ of RA-25298 (2.5uM) or an equivalent volume of DMSO. 24 hours post infection supernatants were collected and infectious virus was quantified using a plaque assay before repeating the experiment. Once the titers of RA-25298 treated replicates had reached the same titers as DMSO three times in a row we sequenced the virus from each of the last three passages using a previously described method (*49*).

### Titer Reduction Assay

MRC-5 fibroblasts were infected with alphaviruses at an MOI of 0.1 for one hour and treated with RA-25298 in 12 point 4 fold dose response from 40uM. Infectious virus in the supernatants were quantified by plaque assay as previously described ((*50*).

### Virus Production and Analysis

Viral RNA was in vitro transcribed from infectious clones as previously described. Mutations were produced in infectious clones of using previously described bubble PCR methods. To produce virus stocks RNA was electroporated in BHK-21 cells and incubated for 24-36 hours depending on the virus. Virus stocks were quantified via plaque assay as described above.

### Protein Purification

CHIKV nsP2 sequence from 181/25 was cloned into a pET28 vector with a N-terminal TwinStrep 6-His SUMO tag and expressed in Rosetta (Novagen) 2 E.coli using previously described autoinduction methods for 48h at 15°C (1). E.coli were lysed in a sodium phosphate buffer (50mM NaHPO_4_ pH7.8, 20% Glycerol, 500mM NaCl, 1mM TCEP) via sonication, nucleic acids were precipitated using polyethylenamine, and the lysate was clarified via centrifugation at 70,000g for 30 minutes. Protein from the clarified lysate was precipitated using ammonium sulfate to 40% saturation and precipitated proteins including nsP2 were pelleted via centrifugation at 70,000g for 30 minutes. Ammonium sulfate was resuspended in sodium phosphate buffer and run over a Streptactin-XT column (Cyvita) and washed with 100 column volumes of buffer prior to elution via on column cleavage by sumo protease ULP1. Elutions were concentrated and dialyzed into a HEPES buffer (25mM HEPES pH 7.4, 250mM NaCl, 20% glycerol, 1mM TCEP).

### Phylogenetic Analysis

Sequences of 31 different alphaviruses were acquired from NCBI and aligned using clustal Omega. To visualize the phylogenetic tree the sequences were analyzed using Geneious to produce a phylogenetic tree. Three groups were identified based on their distance from the center.

### EEEV Replicon Assay

An EEEV strain V105-00210 (KP282670.1) replicon plasmid (pEEEV-nLucRep) was assembled in a pTwist vector from synthesized DNA fragments. EEEV strain V105 was isolated in 2005 from the brain of a fatal human case (*51*). Downstream of an SP6 promoter, the replicon genome encodes the EEEV V105 nonstructural genes and the nanoluciferase gene downstream of the EEEV 26S promoter in place of the virus structural protein genes. Specific mutations in the EEEV V105 nsP2 gene were introduced into the replicon plasmid through site-directed mutagenesis using specific primers and the QuikChange II XL site-directed mutagenesis kit (Agilent Technologies) as previously described (*52*). Briefly, capped viral replicon RNA was generated by in vitro transcription of Not I linearized replicon plasmid DNA templates using the mMessage mMachine SP6 transcription kit (Invitrogen). 10 ng of replicon RNA was transfected into BHK-21 cells using Lipofectamine MessengerMAX reagent according to the manufacturer’s instructions (Invitrogen). At 18-24 h post-transfection, luciferase activity in cells was measured using the Nano-Glo Luciferase Assay System (Promega). Luminescence was measured in CELLSTAR Chimney Well 96 Flat Black plates using a Tecan Infinite M Plex Plate Reader.

### Time of Addition

MRC-5 fibroblasts were infected at an MOI of 3 for 1 hour prior to washing with PBS and DMEM. Following infection RA-25298 was added at 0,1,2,4,6,and 12 hour intervals at an EC90 as calculated from previously run CHIKV nLuc antiviral assays. Supernatant and cells were collected at 24 hours post infection and virus was quantified via plaque assay, protein was measured using western blot for nsP2 (Invitrogen-HL1432) and capsid (Invitrogen.

### Pharmacokinetics

For co-dosing studies, male C57BL/6 mice (n = 3, 6−8 weeks, 20−30 g; Vital River) were dosed by i.v (10 mg/Kg), p.o. (30 mg/Kg) and i.p. (30 mg/Kg) administration of RA-25298 and RA-188293 as a volume of a 2 mg/mL (i.v.) and 3 mg/mL (p.o. and i.p.) solution respectively in DMSO/PEG-400/water (v/v/v, 5:50:45). 1-ABT (100 mg/kg p.o.) was administered as 10 mL/kg volume of a 10 mg/mL solution in saline as a 2 h pretreatment. Blood (0.03 mL) was collected from the dorsal metatarsal vein at 0.08, 0.25, 0.5, 1, 2, 4, 8, and 24 h (for RA-25298) and 0.25, 0.5, 1, and 3 h (for RA-188293) time points and the concentration of the samples in plasma was determined by LC-MS/MS analysis. Concentrations of both the compounds in the plasma samples were determined using an AB API 5500+ LC-MS/MS instrument fitted with an Agilent EC-C18 column (2.7 μm, 3 x 30 mm) using a mobile phase of 5−95% MeCN in H2O with 0.1% formic acid. Pharmacokinetic parameters were calculated from the mean plasma concentration versus time by a non-compartmental model using WinNonlin 8.3 (Phoenix) to determine Cmax, AUClast, t1/2, CLs.

### Plasma protein binding

Frozen plasma that had been stored at −80°C was thawed in a 37°C water bath. Working solutions of test compound RA-25298 and the control ketoconazole were prepared in DMSO at 200 μM and then spiked into plasma. The final concentration of the compounds was RA-25298 μM with 0.5% DMSO. The dialysis membranes were soaked in ultrapure water for 60 min to separate strips, in 20% ethanol for 20 min, and finally in dialysis buffer for 20 min. Each dialysis cell was filled with 150 μL of plasma sample and dialyzed against equal volume of PBS buffer. The assay was performed in duplicate. The dialysis plate was sealed and incubated at 37°C with 5% CO2 at 100 rpm for 6 h. 50 μL of samples from both buffer and plasma chambers were transferred to a 96-well plate. 50 μL of plasma was added to each buffer sample and supplemented with an equal volume of PBS. A 400 μL of quench solution (200 nM labetalol, 100 nM tolbutamide, and 100 nM ketoprofen in acetonitrile) was added to precipitate protein and release compounds. Samples were vortexed for 2 min and centrifuged for 30 min at 3,220 g. 100 µL of the supernatant was diluted with 100 µL of ultra-pure H2O, and the mixture was used for LC-MS/MS analysis.

### In Vivo Efficacy

Six week old male C57BL/6 mice were treated with 100mg/kg 1-ABT (dissolved in saline) 2hrs prior to treatment with 30mg/kg of RA-0188293 or vehicle (DMSO/PEG400/water-5/50/45) and then infected in the left rear foot pad with 1000 PFU of the Sri Lanka-15649 strain of CHIKV. Every 24 hours mice were treated with 1-ABT and every 12 hours with RA-0188293. Dosing was staggered so that each day the first dose of 188293 occurred at 2 hours post 1-ABT treatment. For 5 and 7 DPI experiments, blood was collected at 3DPI via cheek bleed to assess viremia. At the determined time point, mice were euthanized by an overdose of isoflurane (Baxter), blood collected by cardiac puncture, organs and/or feet collected for downstream processing. For pathology studies, mice were euthanized by an overdose of isoflurane (Baxter), blood collected by cardiac puncture and the mouse perfused with 4% PFA followed by submersion in 4% PFA for 1 week. All mouse studies were conducted under protocols approved by the Institutional Animal Care and Use Committee (IACUC) at the University of North Carolina at Chapel Hill.

Infectious viral loads were measured by plaque assay. Tissues (feet, quads, and spleens) were homogenized in 1mL of media (DMEM + 5% FBS + 1% L-glutamine) 3 consecutive times using settings 6.0m/s for 30 sec on a Beadblaster homogenizer (Benchmark Scientific). Centrifugation at maximum speed for 1 minute removed tissue debris. A 50μL aliquot of the clarified homogenate was added to 450μL of dilution media (DMEM + 5% FBS + 1% L-glutamine media) and subsequently serially diluted to create ten-fold serial dilutions (10^-1^ to 10^-6^). Approximately 200uL of each dilution was pipetted onto previously plated Vero81 cells and incubated at 37°C for 2 days with 2mL of media+carboxymethylcellulose (50:50 mixture of 2X alpha MEM containing 6% FBS + 2% penicillin/streptomycin + 2% L-glutamine + 2% HEPES and 2.5% carboxymethylcellulose). After incubation for 2 days an equal volume of 4% paraformaldehyde was added to each well and the cells were allowed to fix overnight. The fixative was removed, wells were rinsed with water to remove residual overlay and 0.25% crystal violet was added to each well. Visible plaques were counted and averaged between two technical replicate wells and used to calculate plaque forming units (pfu) per mL of tissue. The limit of detection (LOD) for the assay was determined to be 25 pfu/tissue, and samples that yielded no plaques were assigned a value of 12.5, half of the LOD.

### Deuterium exchange

Labeling buffer was prepared by diluting 20 X Tris-HCl, MgCl in H_2_O in D_2_O (99.9%). 3 μL of the sample were added to 57 μL of labeling buffer for a final labeling concentration of 90.16%. Deuterium labeling was carried out for 1 and 10 min at 20°C. During HDXMS experiments protein samples were stored at 0°C and stability was assessed by staggering technical replicates. After labeling, equivalent volumes of labeling reaction and prechilled quench solution (1.5 M GdnHCl, 0.25 M TCEP) was added to bring the reaction to pH 2.5. Reaction conditions are summarized in Supplementary Table 1.

#### Mass spectrometry and peptide identification

Approximately 40-50 μmol of the sample were loaded onto a BEH pepsin column (2.1 × 30 mm) (Waters, Milford, MA) in 0.1% formic acid at 75 μL/min. Proteolyzed peptides were trapped in a C18 trap column (ACQUITY BEH C18 VanGuard Pre-column, 1.7 µM, Waters, Milford, MA). Peptides were eluted in an acetonitrile gradient (8–40%) in 0.1% formic acid on a reverse phase C18 column (AQUITY UPLC BEH C18 Column, Waters, Milford, MA) at 40 μL/min. All fluidics were controlled by nanoACQUITY Binary Solvent Manager (Waters, Milford, MA). Electrospray ionization mode was utilized, and ionized peptides were sprayed onto an SYNAPT XS mass spectrometer (Waters, Milford, MA) acquired in HDMS^E^ Mode. Ion mobility settings of 600 m/s wave velocity and 197 m/s, transfer wave velocity were used with collision energies of 4 and 2 V were used for trap and transfer, respectively. High collision energy was ramped from 20 to 45 V while a 25 V cone voltage was used to obtain mass spectra ranging from 50 to 2000 Da (10 min) in positive ion mode. A flow rate of 5 µL/min was used to inject 100 fmol/μL of [Glu^1^]-fibrinopeptide B ([Glu^1^]-Fib) as lockspray reference mass.

Peptides of CHIKV nsp2 were identified through independent searches of mass spectra from the undeuterated samples in two steps. Peptides were identified from a database containing the amino acid sequence of nsp2 using PROTEIN LYNX GLOBAL SERVER version 3.0 (Waters, Milford, MA) software in HDMS^E^ mode for non-specific protease cleavage. Search parameters in PLGS were set to ‘no fixed or variable modifier reagents’.

Deuterium exchange was quantitated using DynamX v3.0 (Waters, Milford, MA) with cutoff filters of: minimum intensity = 2000, minimum peptide length = 4, maximum peptide length = 25, and precursor ion error tolerance <10 ppm. Four undeuterated replicates were collected for CHIKV nsp2, and the final peptide list includes only peptides that fulfilled the above-described criteria and were identified independently in at least 3 of the 4 undeuterated samples.

### Hydrogen deuterium exchange analysis

The average number of deuterons exchanged in each peptide was calculated by subtracting the centroid mass of the undeuterated reference spectra from each deuterated spectra. Peptides were independently analyzed for quality across technical replicates. Relative deuterium exchange and difference plots were generated by DynamX v3.0. The data for the mass spectra were acquired from DynamX v3.0. Relative deuterium exchange plots are reported as RFU which is the ratio of exchanged deuterons to possible exchange deuterons. Deuteros (53) was used to generate the butterfly plots and pymol scripts for significantly different peptides using hybrid significance testing (p<0.05) (54). The mass spectrometry proteomics data will be deposited to the ProteomeXchange Consortium via the PRIDE partner repository.

### Chemical Synthesis

Compounds were synthesized by the Structural Genomics Consortium at the University of North Carolina Chapel Hill as previously described (*41*).

## Supporting information

Combined Supplemental Figures

