## Supplementary material for "Identification of a Broadly Acting Inhibitor of the Alphavirus Non-Structural Protein 2 Helicase": Combined Supplemental Figures

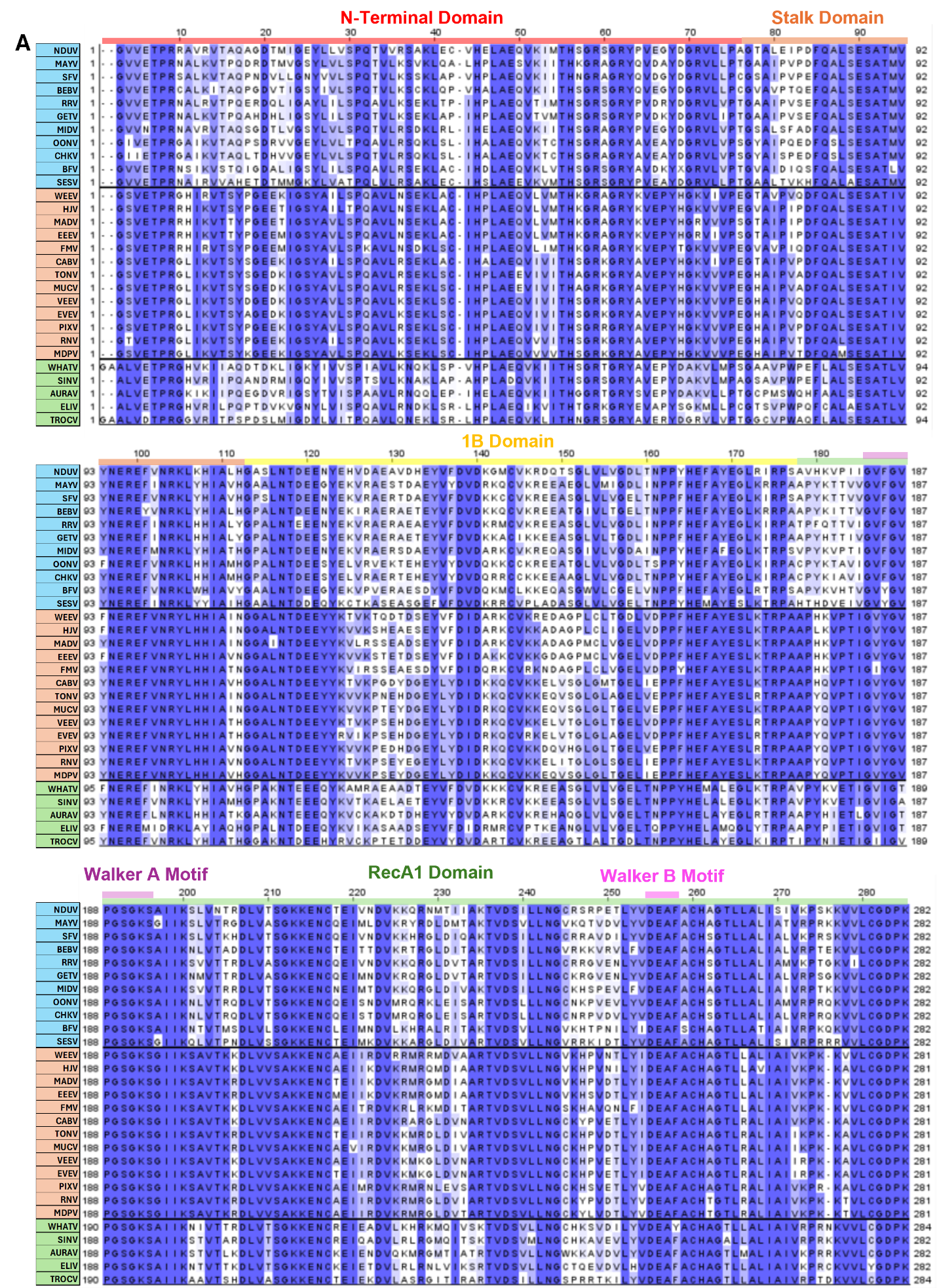

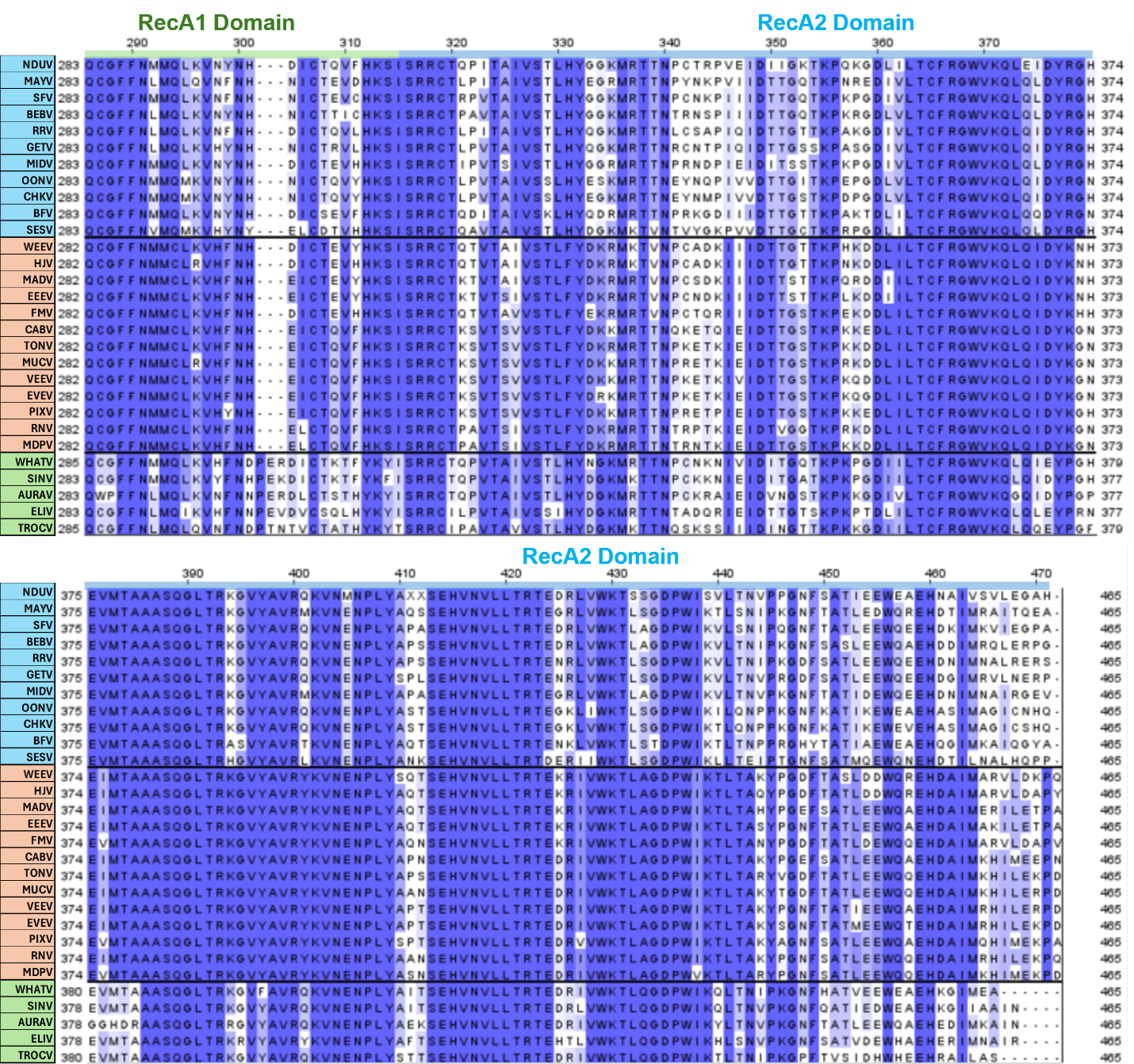
**Supplemental Figure 1. Alignment of alphavirus nsP2 helicase domain sequences.** (A) The first 465 amino acids of 29 different alphavirus nsP2 sequences were aligned using Clustal Omega. Different domains and motifs are denoted with colored bars. Shading indicates the degree of sequence conservation, with more conserved regions in darker blue and less conserved regions in white.

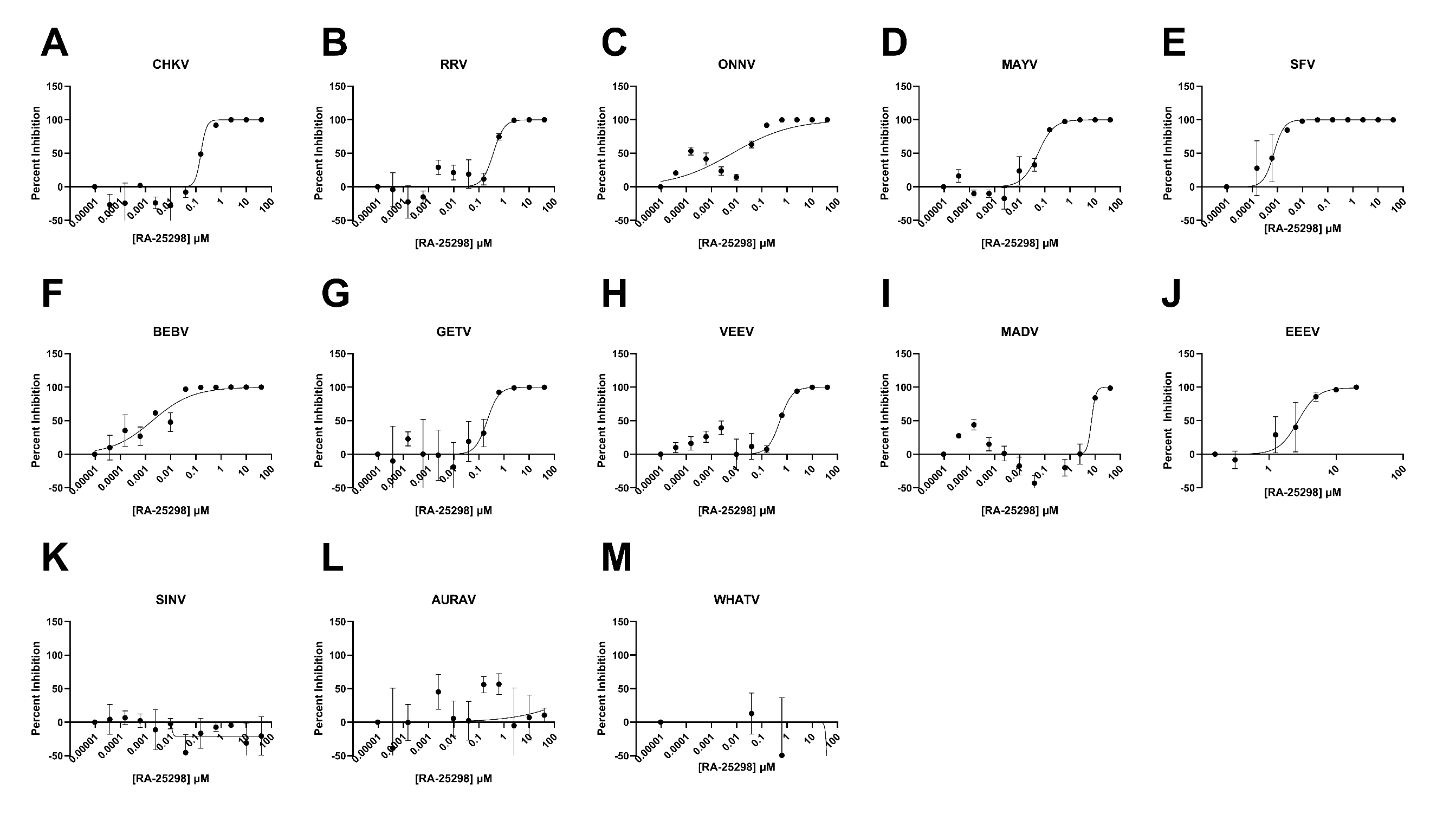
**Supplemental** **Figure 2. Hill-slope curves of dose response inhibition by RA-25298 on different alphaviruses.** (A-M) Percent inhibition of replication of a diverse group of alphaviruses by RA-25298 was calculated by comparing all values to the lowest tested concentration. Values were fitted to a 4 parameter hill-slope equation to determine EC_50_ values.

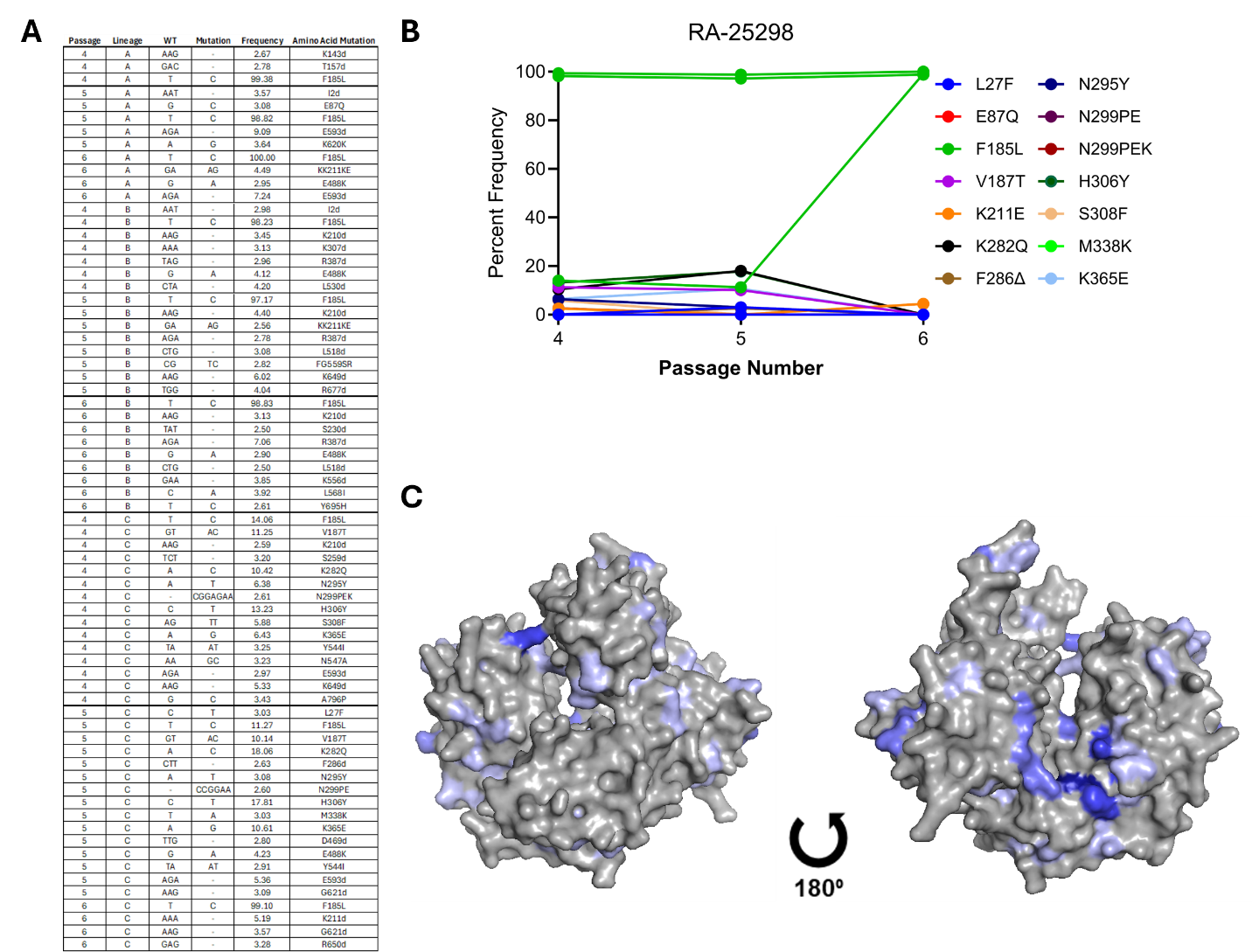
**Supplemental Figure 3. Passaging chikungunya virus in the presence of RA-25298 identifies multiple mutations in nsP2.** MRC-5 fibroblasts were infected with CHIKV in the presence of 1.25uM RA-25298 at an MOI=0.1. 24 hours post infection virus was collected passaged on fresh cells. The process was repeated six times, after which viral RNA was extracted from supernatants, and the nsP2 region was sequenced. Mutations that occurred at a frequency > 2.5% are shown in (A). The frequency of the 14 most common mutations in passages 4, 5 or 6 is shown in (B). (C) The mutations shown (B) were mapped onto the structure of the CHIKV nsP2 helicase domain. Darker blue shading corresponds to higher mutation frequency.

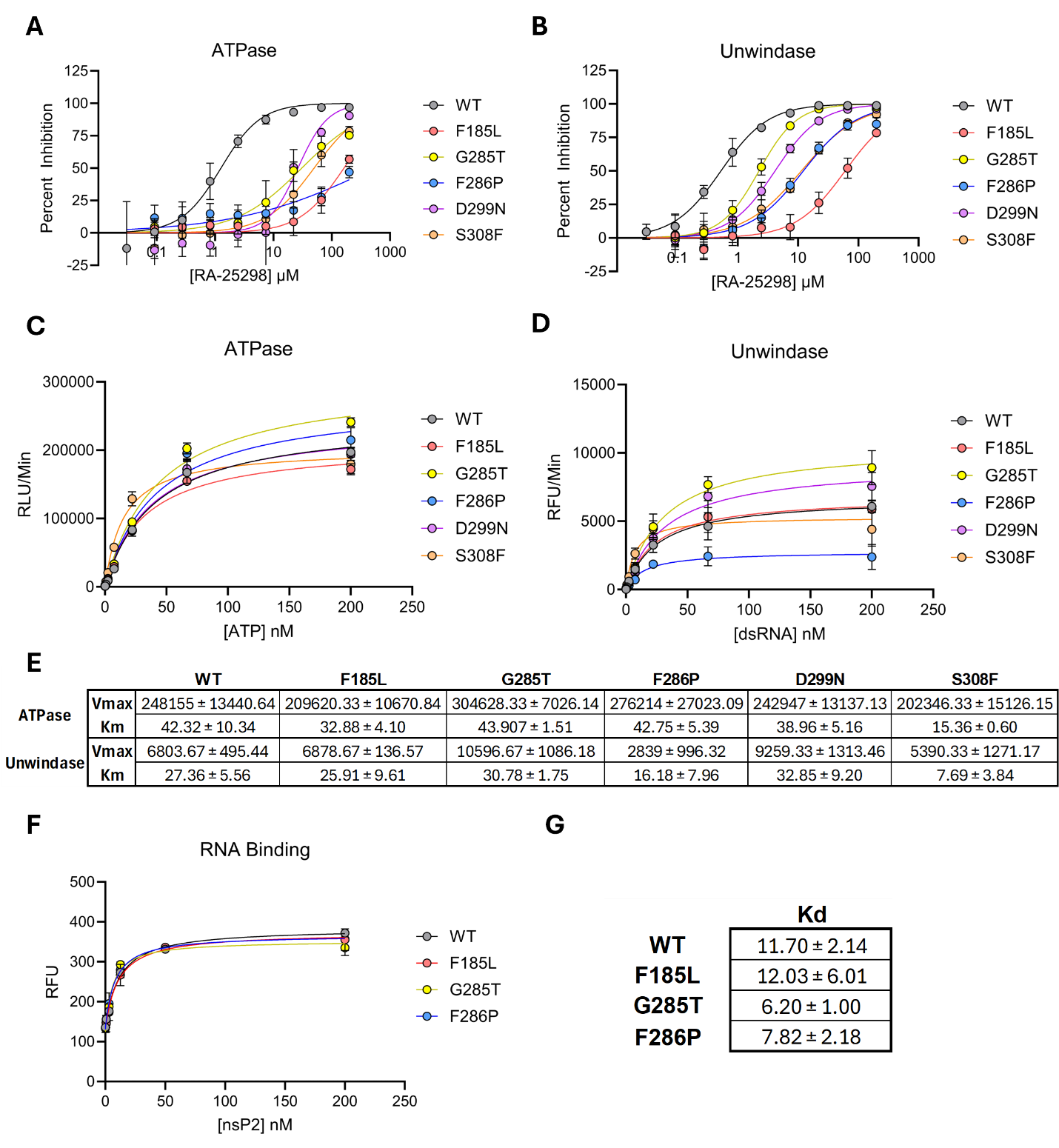
**Supplemental Figure 4. Enzymatic analysis of resistance mutants.** (**A,B**) Full length nsP2 protein containing mutations conferring resistance to RA-25298 in antiviral assays were purified and tested in enzymatic assays measuring RA-25298 inhibition of (**A**) ATPase and (**B**) unwindase activity. The ATPase and unwindase activity of each mutant was also measured in the presence of increasing concentrations of (**C**) ATP or (**D**) RNA substrate (**E**) Michaelis constants for the indicated mutants. (**F**) RNA binding was measured for the indicated mutants using fluorescence polarization and the data used to calculate Kd (**G**).

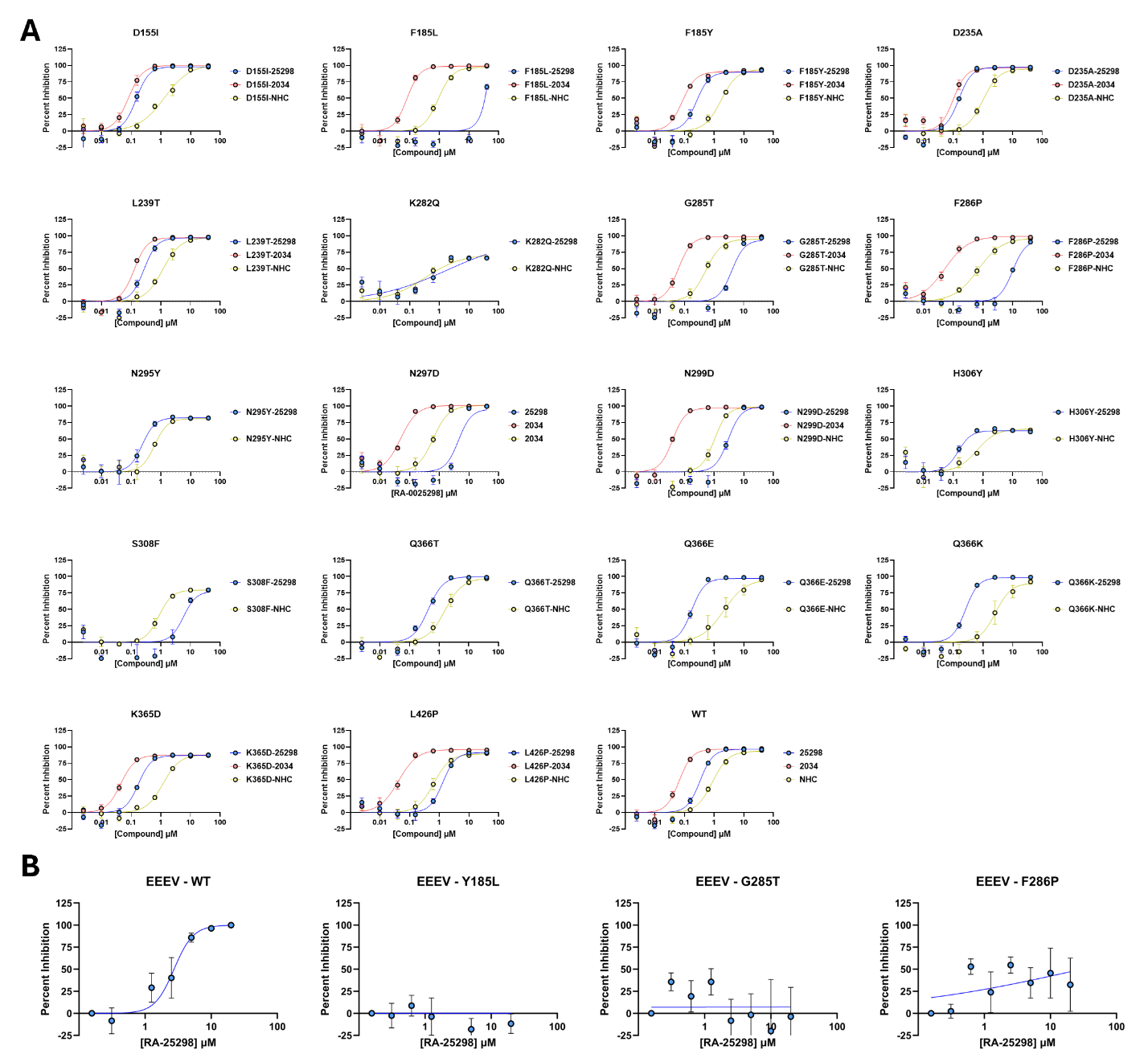
**Supplemental Figure 5. Antiviral dose response for all mutants tested against RA-25298.** (A) Antiviral activity of RA-25298, the protease inhibitor RA-000234, and the nsP4 polymerase inhibitor NHC against a series of CHIKV nsP2 point mutants generated in a reporter virus expressing nanoluciferase. MRC-5 fibroblasts were treated with an 8-point 4 fold dose response of compound starting at 40µM 1 hour prior to infection with virus. Luciferase production was measured 6 hours post infection and the percent inhibition was determined by comparing treated wells to DMSO control wells. (B) Antiviral activity of RA-25298 against orthologous mutants in EEEV was assessed using a similar protocol as A. Data points were fitted to 3 parameter hill slope equations to determine EC_50_.

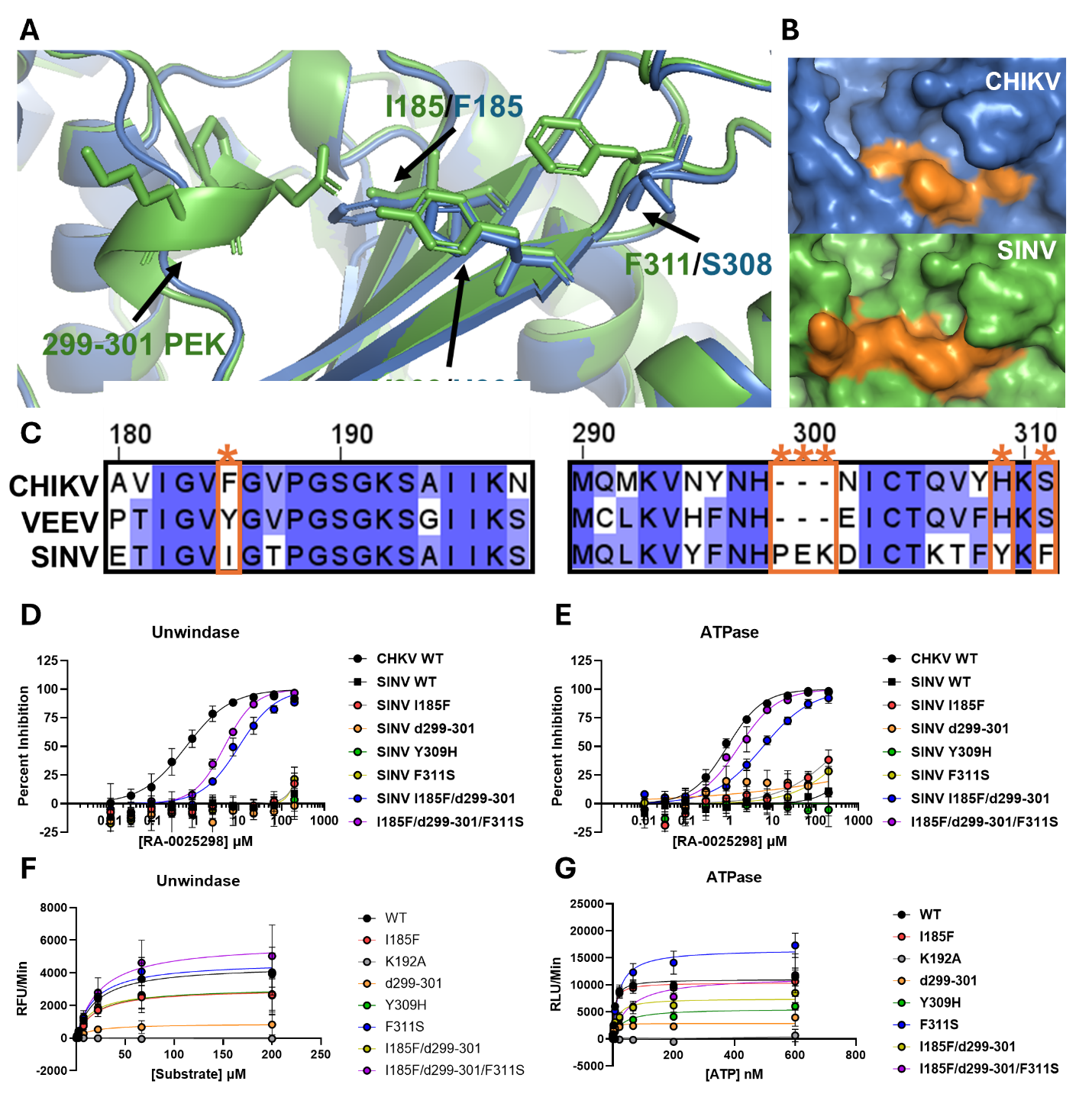

**Supplemental Figure 6. Mutation of pocket that differs between SINV and other alphaviruses restores susceptibility to RA-25298.** (A-C) Differences in amino acid composition and structure of pocket on nsP2 where resistance mutations arise. (D) SINV mutant protein was assessed for ability of RA-25298 to inhibit unwindase and (E) ATPase activity. (F-

**A**

G) Impact of mutations on SINV nsP2 unwindase and ATPase activity as determined by Michaelis-Menten kinetics.

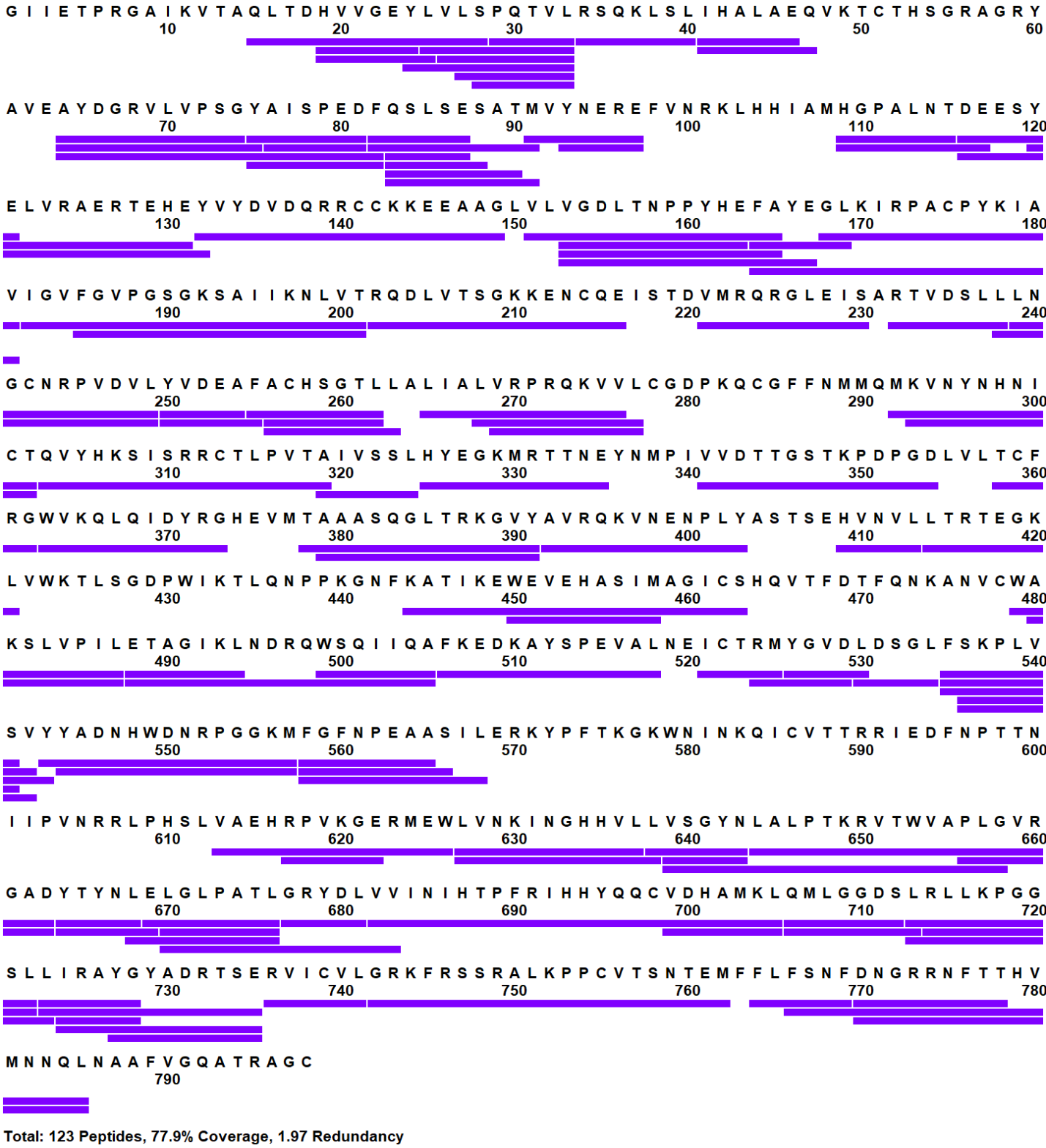

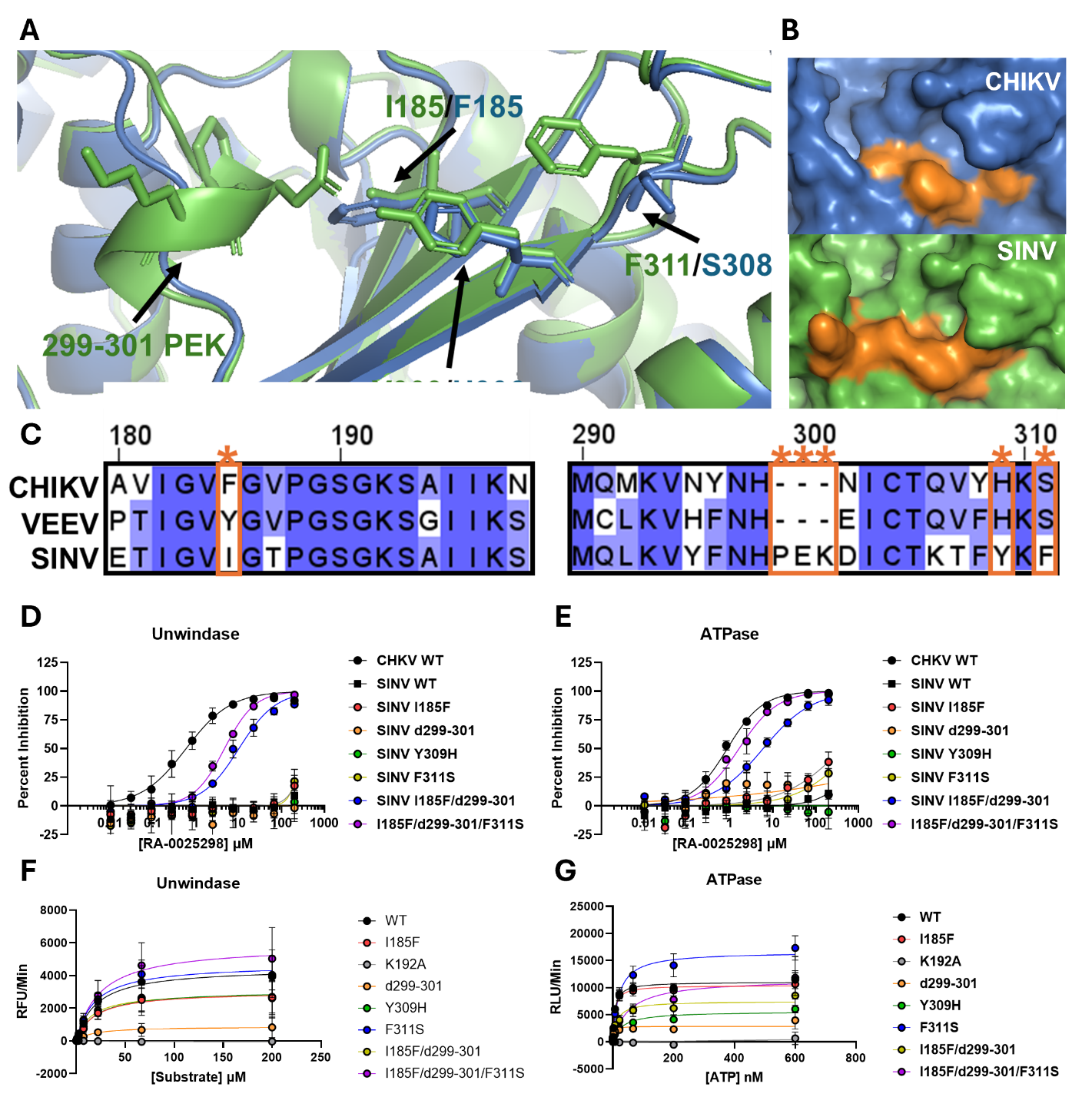
**Supplemental Figure 7.** (A) Sequence coverage by pepsin fragmentation of CHIKV nsp2 in triplicate.

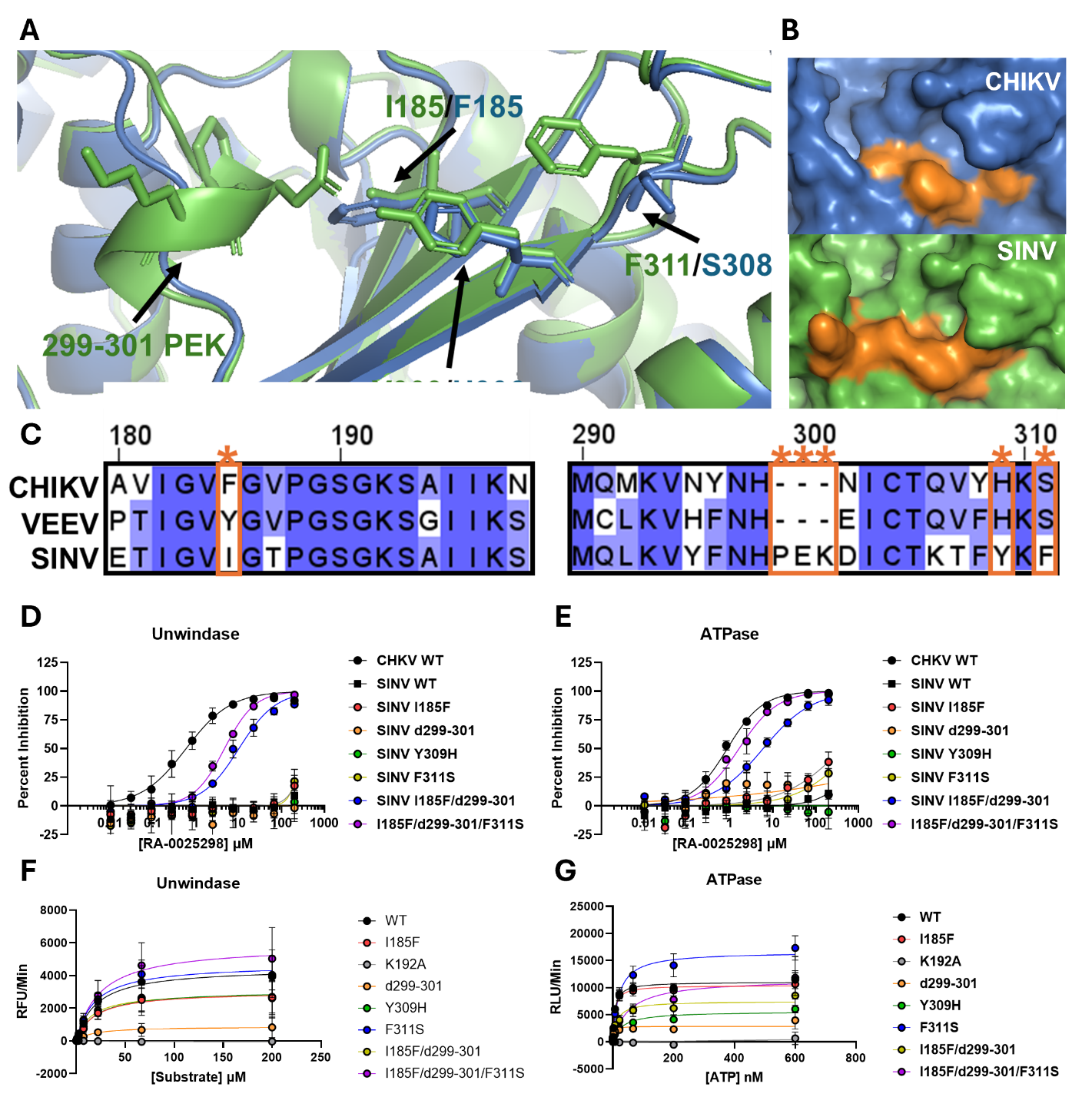

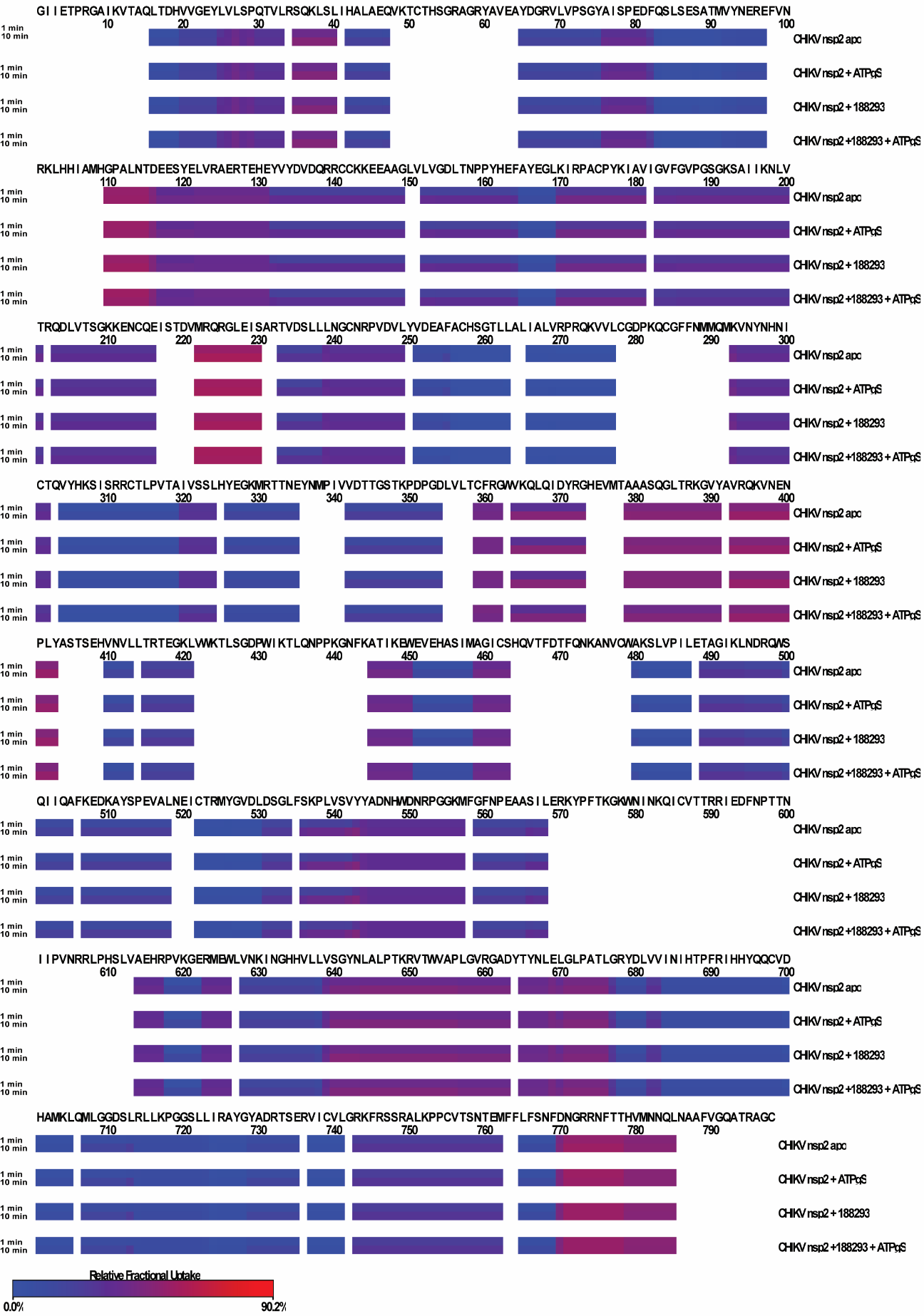
**Supplemental Figure 8**. (A) Deuterium exchange heatmap (D_ex_ = 1 and 10 min) of CHIKV nsp2 peptides from apo, ATPγS, and 188293 (SGC-NSP2hel-1) treated samples.

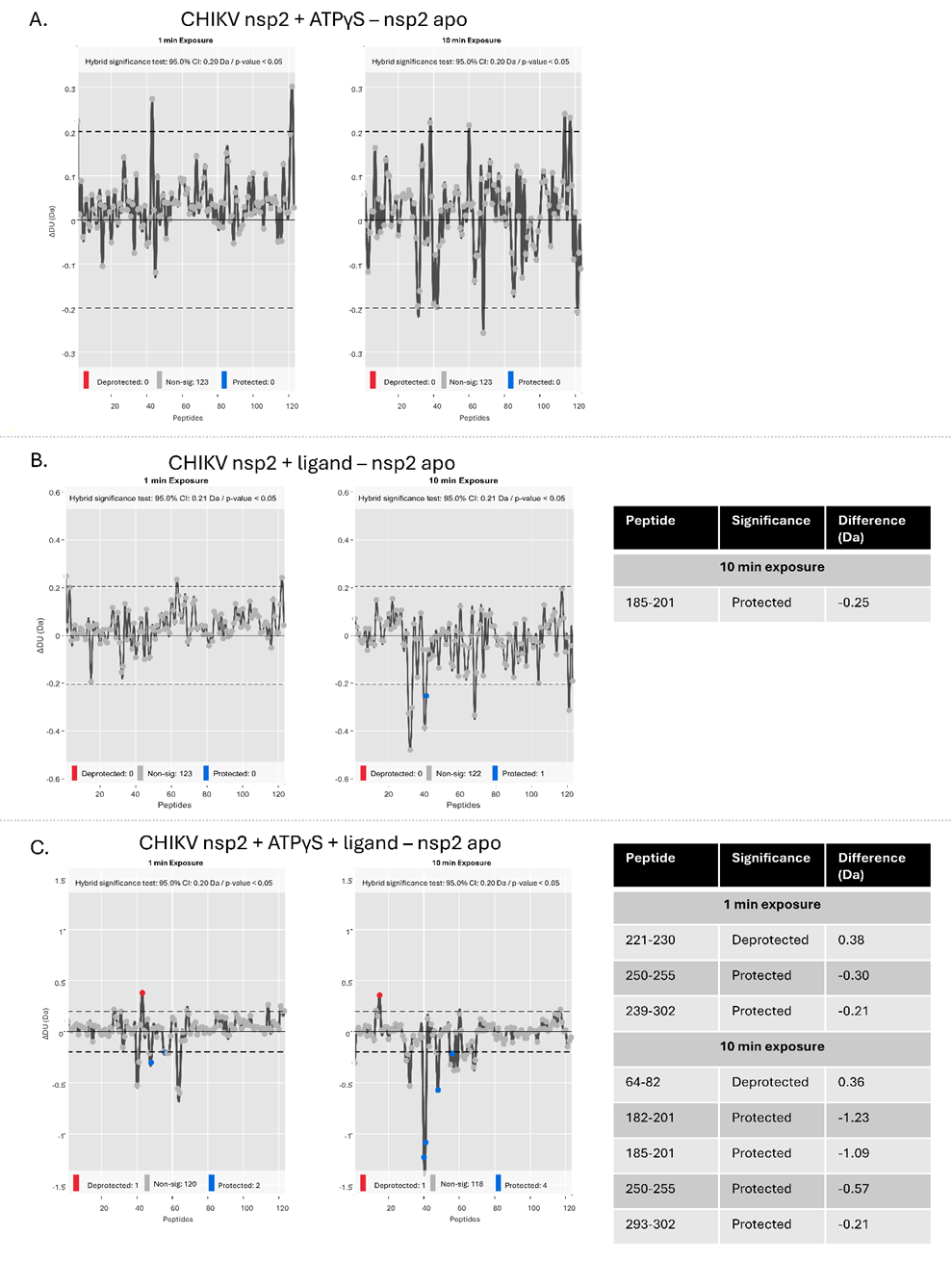

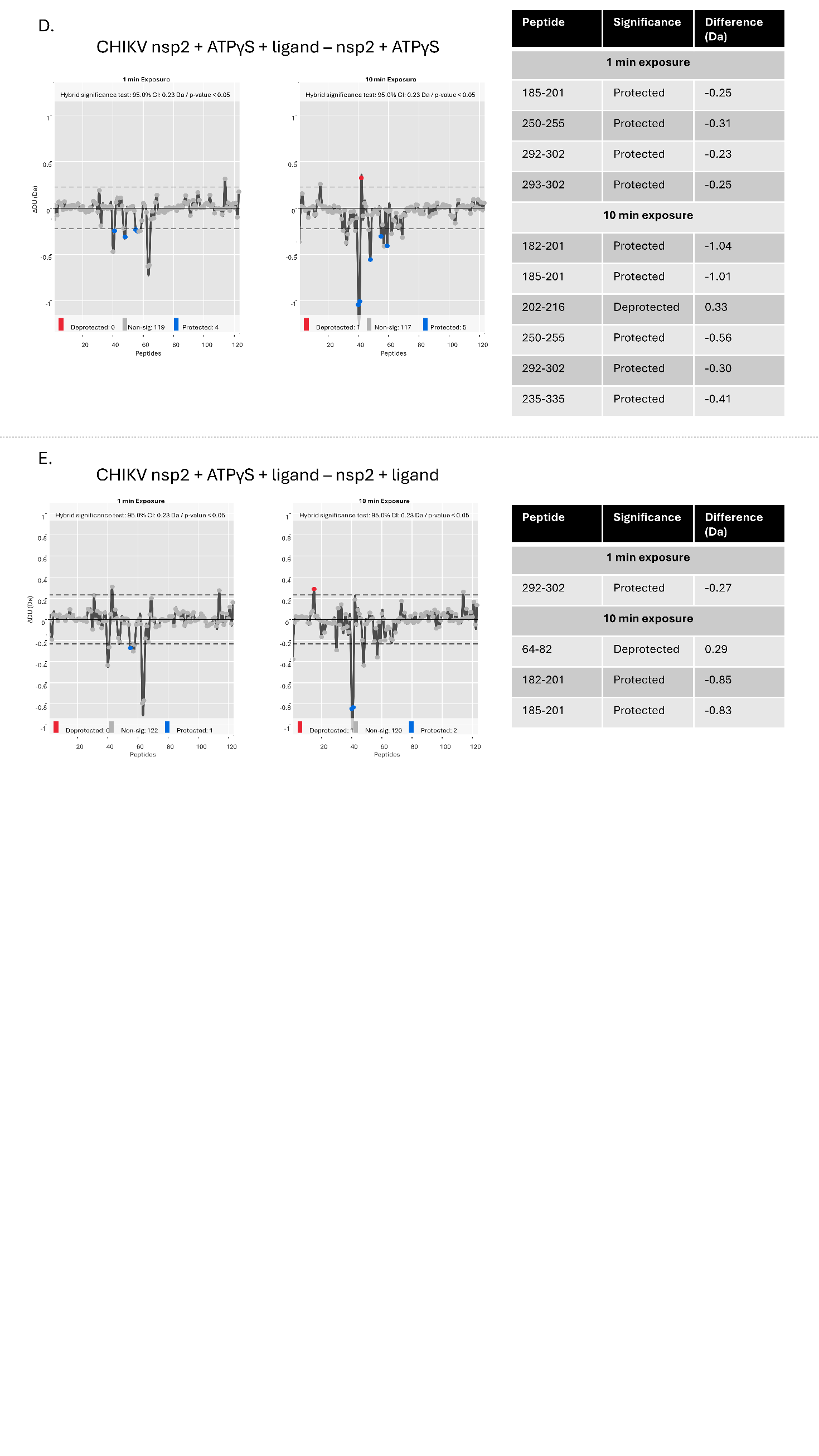
**Supplemental Figure 9.** Differences in deuterium exchange mapped at peptide resolution from N to C terminus for CHIKV nsp2 protein for D_ex_ = 1 and 10 min for (**A)** nsp2 incubated with ATPγS minus apo nsp2, (**B)** nsp2 incubated with ligand minus apo nsp2, (**C)** nsp2 incubated with ATPγS and ligand minus apo nsp2, (**D)** nsp2 incubated with ATPγS and ligand minus nsp2 incubated with ATPγS, and (**E)** nsp2 incubated with ATPγS and ligand minus nsp2 incubated with ligand. Select peptides showing significant differences are designated in corresponding tables.

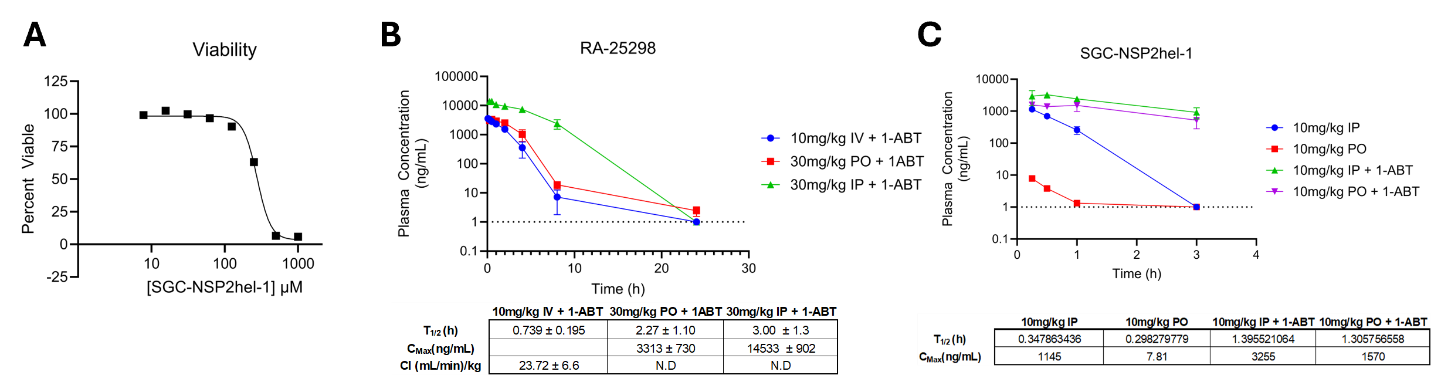
**Supplemental Figure 10. Viability and pharmacokinetic analysis of SGC-NSP2hel-1.** (A) Viability assays were performed on MRC-5 cells on an 8-point 4 fold dose response over 72 hours with fresh compound added every 24 hours. (B) Pharmacokinetic analysis of RA-25298 and (C) SGC-NSP2hel-1was performed using 10 or 30 mg/kg dose of compound with or without pre-treatment with p450 inhibitor 1-ABT in C57J/B6 mice with IV, PO, and IP delivery. Plasma concentration was determined at different time points to determine T_1/2_, C_Max_, and C_I_ were determined from blood plasma concentration.

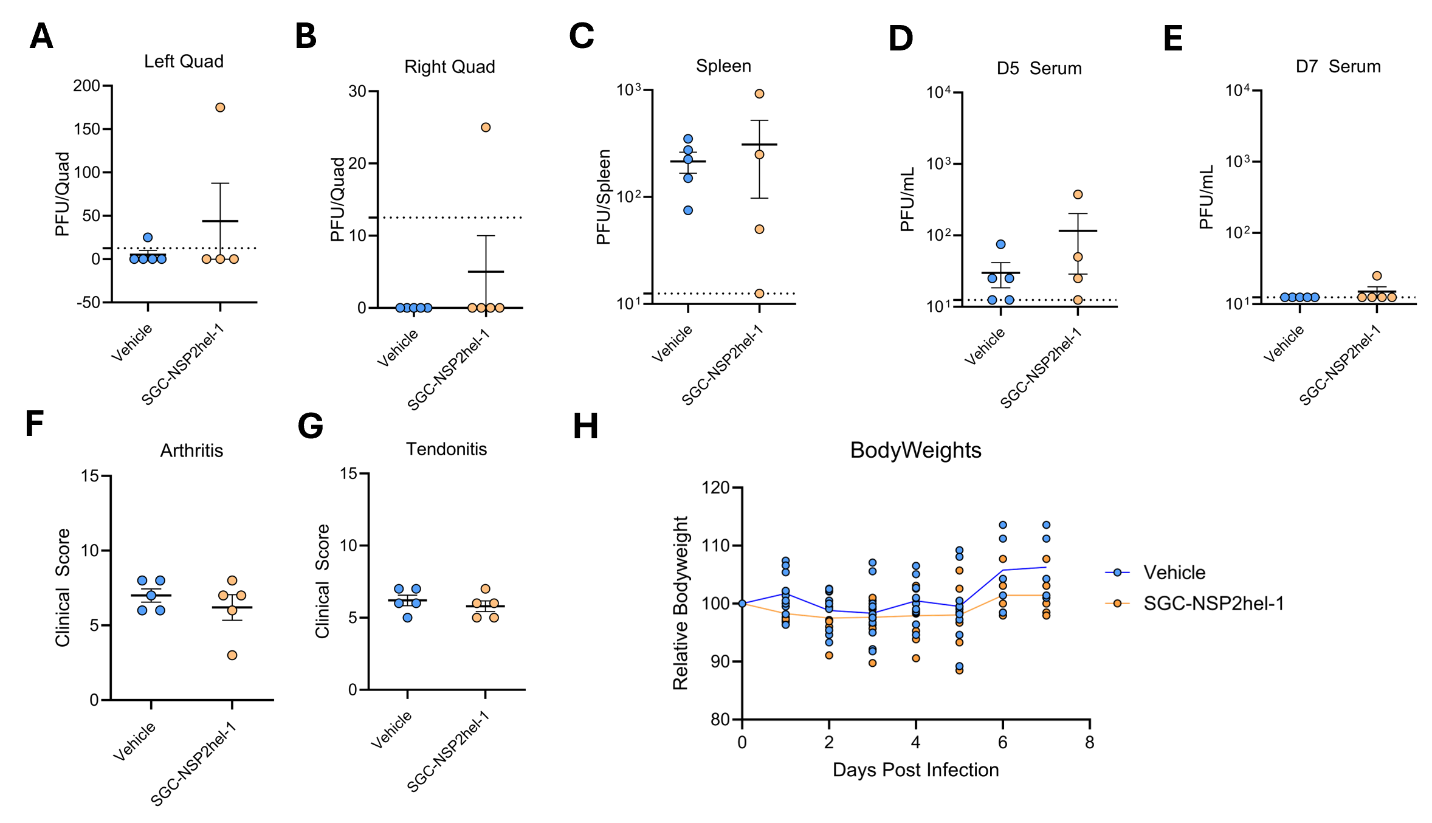
**Supplemental Figure 11. Viral titers and pathology scores from in vivo efficacy experiment with 30 mg/kg SGC-NSP2hel-1.** (A-E) Viral titers in the indicated tissues were measured on the indicated day after infection via plaque assay. (F,G) Clinical pathology scores for tendonitis and arthritis were determined from H&E stains. (H) Mice were weighted daily as a measure of animal health.

| **HDX reaction details** | Labeling buffer: 40 mM Tris-HCl, 20 mM MgCl pH 7.49 prepared in 99.9% D_2_O to a final concentration of 94.9% D_2_O  Quench buffer: 1.5 M Guandinium hydrochloride, 0.25 M TCEP  CHIKV nsp2 apo HDXMS reaction: 57 μL Labeling buffer added to 3 μL CHIKV nsp2 sample for deuterium exchange at 20°C and quenched with 60 μL of quench buffer by bringing reaction to pH 2.5 at 0°C.  CHIKV nsp2 + ligand HDXMS reaction: Mixed together 1 μL of ligand with 73 μL of nsp2 and incubated. Added 57 μL Labeling buffer to 3 μL CHIKV nsp2 + ligand sample for deuterium exchange at 20°C and quenched with 60 μL of quench buffer by bringing reaction to pH 2.5 at 0°C.  CHIKV nsp2 + ATPγS HDXMS reaction: Mixed together 1 μL of ATPγS with 73 μL of nsp2 and incubated. Added 57 μL Labeling buffer to 3 μL CHIKV nsp2 + ATPγS sample for deuterium exchange at 20°C and quenched with 60 μL of quench buffer by bringing reaction to pH 2.5 at 0°C.  CHIKV nsp2 + ligand + ATPγS HDXMS reaction: Mixed 1 μL of ligand and 1 μL of ATPγS with 73 μL of nsp2 and incubated. Added 57 μL Labeling buffer to 3 μL CHIKV nsp2 + ligand + ATPγS for deuterium exchange at 20°C and quenched with 60 μL of buffer bringing reaction to pH 2.5 at 0°C. |
| --- | --- |
| **Incubation** | ≥20 minutes at 20°C |
| **Concentration** | CHIKV nsp2: 1.4 mg/mL (15.6 μM)  ATPγS: 4.56 mM  Ligand: 4.56 mM |
| **HDXMS un-deuterated controls** | 57 μL of 40 mM Tris-HCl, 20 mM MgCl, pH 7.49 was added to 3 μL of sample and 60 μL of quench buffer was added by bringing reaction to pH 2.5 at 0°C. |
| **HDXMS time course** **(min)** | 1, 10 |
| **# of peptides** | 123 |
| **Coverage (%)** | 77.9 |
| **Peptide redundancy** | 1.97 |

**Supplementary Table 1.** Table describing HDX-MS experimental conditions.
